# Hybrid *Drosophila* cell lines show selective genome instabilities

**DOI:** 10.64898/2026.10.27.762186

**Authors:** Jie Hua, Andrea Lukacs, Tatyana G. Kahn, Jacob Cooper, Andreas W. Thomae, Yuri B. Schwartz, Nitin Phadnis, Rupam Choudhury, Ignasi Forné, Axel Imhof

**Affiliations:** Chair of Molecular Biology, Biomedical Center (BMC), LMU Medizin, Ludwig-Maximilians-Universität München, Planegg-Martinsried, Germany; Department of Molecular Biology, Umeå University, Umeå, Sweden; School of Biological Sciences, University of Utah, Salt Lake City, UT, USA; Core Facility Bioimaging, Biomedical Center (BMC), LMU Medizin, Planegg-Martinsried, Germany; Protein Analysis Unit, Biomedical Center (BMC), LMU Medizin, Ludwig-Maximilians-Universität München, Planegg-Martinsried Germany

**Author notes:** **Email addresses:** Jie Hua, Andrea Lukacs, Tatyana G. Kahn, Jacob Cooper, Andreas W. Thomae, Yuri B. Schwartz, Nitin Phadnis, Rupam Choudhury, Ignasi Forné, Axel Imhof.

**Keywords:** Hybrid incompatibility, Drosophila, Speciation, Genome evolution, Dosage compensation, Nucleolar dominance, Ribosomal DNA, Proteomics

## Abstract

**Background:** Post-zygotic reproductive isolation is thought to be caused by genomic conflict. However, the exact nature of these detrimental incompatibilities is largely unknown. To identify genomic regions that became incompatible during the formation of species and are therefore prone to get lost in hybrids, we established hybrid cell lines derived from an authentic interspecies cross between *Drosophila melanogaster* and *Drosophila simulans*.

**Results:** All four independently derived lines selectively lost the *D. melanogaster* allele of the hybrid incompatibility gene *Hmr*, confirming its incompatibility with cellular viability. Moreover, each line underwent distinct but directionally biased erosion of additional *D. melanogaster* chromosomal regions, preferentially affecting the X chromosome, chromosome 4, and chromosome 3R. Loss of the *D. melanogaster* X was dosage compensated by upregulation of genes on the hemizygous *D. simulans* X, while *D. melanogaster* rDNA was paradoxically amplified across all lines. Nucleolar dominance was driven by relative rDNA copy number rather than species identity, overturning the prevailing view of fixed species-based dominance. Finally, the measurements of the species-specific proteome revealed species specific differences in the RNA to protein coefficient K_p_ as novel and yet undescribed sources of molecular conflict in hybrids.

**Conclusions:** These findings reveal a hierarchical, constraint-driven mechanism of hybrid genome resolution.

## Background

Interspecific hybridization forces divergent genomes into a shared cellular environment, disrupting co-evolved regulatory systems and generating hybrid incompatibility, a principal barrier to gene flow and a driver of speciation [1,2]. The Bateson–Dobzhansky–Muller (BDM) model predicts that incompatibilities arise from deleterious interactions between alleles that evolved independently in different genetic backgrounds [3]. Although this framework explains how incompatibilities arise, it provides less insight into what happens when incompatible parental genomes are maintained within the same cellular environment. In *Drosophila melanogaster × D. simulans* crosses, post-zygotic lethality is caused by mis-regulation of the rapidly co-evolving chromatin proteins HMR and LHR, which localize to pericentromeric heterochromatin and telomeres, associate with HP1a, and together with centromeric proteins NLP, NPH and the uncharacterized factors BOH1 and BOH2, form the Hybrid Incompatibility Complex (HIC) [4,5]. Although the molecular basis of this system is well characterized, it remains unknown whether the genomic conflict leads to a stochastic genomic deterioration or follows intrinsic constraints toward a limited set of viable, stable configurations.

Evidence from diverse hybrid systems suggests that genome remodeling is non-random: mouse– human somatic hybrids preferentially eliminate one parental genome [6,7]; Xenopus interspecies hybrids eliminate paternal chromosomes via chromothripsis [8]; plant hybrids show uniparental chromosome loss linked to centromere assembly defects [9]; and natural stickleback hybrids undergo rapid, genome-wide purging of one parental genome following tsunami-induced hybridization [10] Centromeres have emerged as a central arena of inter-species conflict, as centromeric satellite DNA evolves rapidly and centromere-associated proteins must co-evolve to maintain function [11,12]. Yet existing studies remain largely descriptive and lack a mechanistic framework for how hybrid genomes resolve conflicts.

To address this gap, we established four independently derived hybrid cell lines from female *D. melanogaster ×* male *D. simulans* crosses and performed integrated long-read genomic, transcriptomic, and proteomic analyses to compare the structural and functional outcomes of genomic conflict across independently derived viable cell lines. Unlike previous somatic hybrid models generated by artificial cell fusion between distantly related species, our lines arise from an authentic interspecies cross in an established genetic model system with fully characterized incompatibility genes providing a tractable platform for identifying the genetic and genomic constraints that shape viable hybrid states.

## Results

### Establishment of D. melanogaster × D. simulans hybrid cell lines

Hybrid cell lines were generated by crossing *D. melanogaster* females carrying Act5C-GAL4 (genotype: y1 w*; P{Act5C-GAL4}25FO1/CyO, y+) with *D. simulans* males carrying UAS-Ras^V12^ in a w501 background, following a modified Simcox protocol [13,14] **(Fig. 1a)**. Of eighteen independently initiated hybrid cultures, only four survived the prolonged establishment period and gave rise to immortalized cell lines after approximately 3–7 months (>10 passages); these were designated HYC1–HYC4. The remaining fourteen cultures progressively ceased proliferation and were lost. Thus, HYC1–HYC4 represent four independently recovered viable outcomes from a stringent hybrid-cell establishment bottleneck. Reciprocal crosses (*D. simulans* mothers × *D. melanogaster* fathers) yielded no viable lines, consistent with the known direction-specificity of hybrid lethality. The *D. melanogaster* parental control (Ras3) was generated from an intraspecific Act5C-GAL4 × UAS-Ras^V12^ cross [15]; the *D. simulans* control (ML82) was the spontaneously immortalized line ML82-19a from the Drosophila Genomics Resource Center (DGRC).

**Fig. 1:**
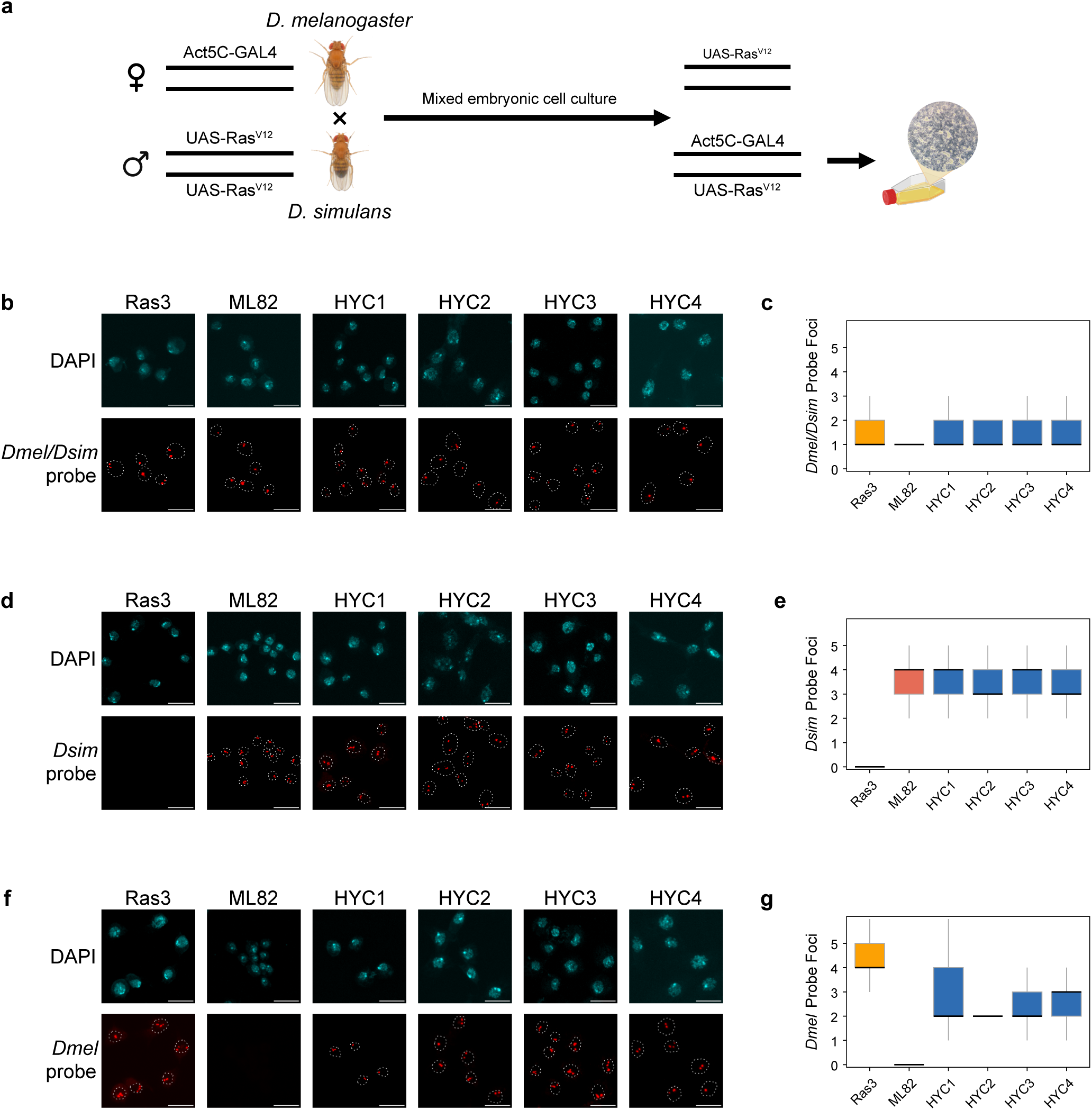
Establishment and validation of Drosophila melanogaster × Drosophila simulans hybrid cell lines. a, Schematic of the crossing strategy and cell-line establishment workflow used to generate HYC1–HYC4 from *D. melanogaster* females carrying Act5C-GAL4 and *D. simulans* males carrying UAS-RasV12. **b,d,f,** Representative DNA fluorescence in situ hybridization images of the parental control lines Ras3 (*D. melanogaster*) and ML82 (*D. simulans*) and the hybrid lines HYC1–HYC4, using probes recognizing sequences shared between the two species (b), *D. simulans*-specific sequences (d) or *D. melanogaster*-specific sequences (f). DNA was counterstained with DAPI. Scale bars, 10□μm. **c,e,g,** Quantification of DNA-FISH foci per nucleus detected with the shared-sequence (c), *D. simulans*-specific (e) and *D. melanogaster*-specific (g) probes. More than 50 nuclei were analyzed per cell line. Box plots show the median and interquartile range.

These cell lines are distinct from all previously described hybrid somatic systems in two critical aspects. First, they are derived from an authentic interspecies cross in a genetically tractable animal model, rather than from artificial polyethylene-glycol-mediated cell fusion. Second, they emerge within a well-characterized incompatibility framework both parental strains carry defined transgenes (Act5C-GAL4 and UAS-Ras^V12^) that drive Ras^V12^-mediated immortalization only in hybrid cells providing unprecedented genetic control over cell line origin. Because immortalization requires expression of UAS-Ras^V12^ (contributed by *D. simulans* males) driven by Act5C-GAL4 (contributed by *D. melanogaster* females), only genuine hybrid cells are selected. This design makes HYC1–HYC4 uniquely suited to study how natural hybrid incompatibilities are resolved at the cellular level.

DNA-FISH using species-specific probes confirmed the presence of both parental genomes in all four hybrid lines, while parental controls (Ras3 and ML82) showed signal only with their respective species-specific probes as well as probe against common sequences **(Fig. 1b,d,f)**. Quantification revealed that whereas the shared-sequence and *D. simulans*-specific probes detected comparable foci across lines, the *D. melanogaster*-specific probe showed a consistent reduction in hybrid lines relative to Ras3, most pronounced in HYC3 **(Fig. 1c,e,g)** a first indication of asymmetric genomic composition, which we subsequently confirmed by long-read sequencing. Because the *D. simulans* X chromosome can only be paternally inherited in female embryos in this cross direction, its presence in all four hybrid lines indicates that the hybrid lines originated from female hybrid embryos **(Fig. 2a)**.

**Fig. 2:**
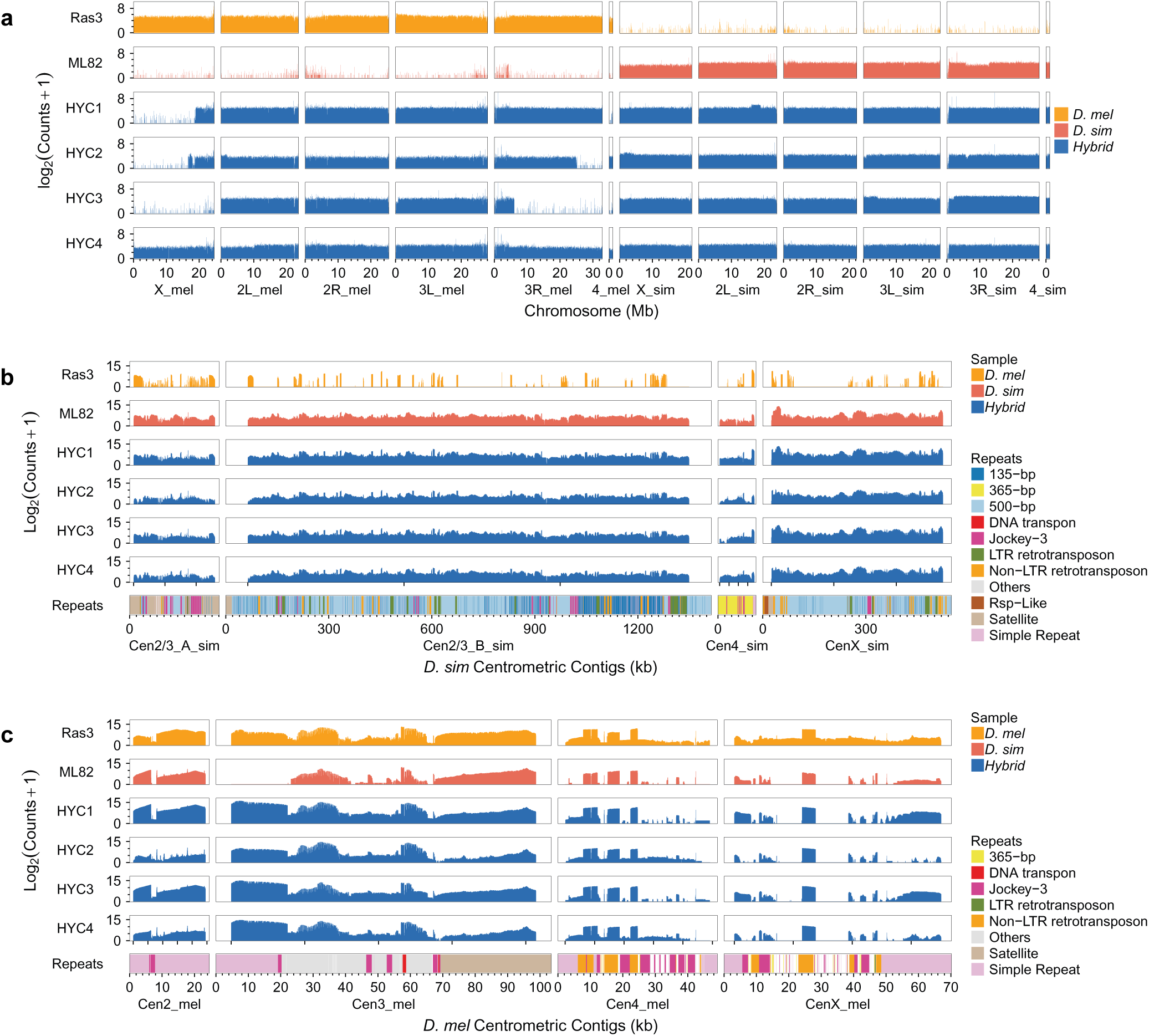
Asymmetric loss of *D. melanogaster* genome content in hybrid cell lines. a, Genome-wide DNA-sequencing read depth across the major chromosomes of Ras3, ML82 and HYC1–HYC4. Read counts were calculated in non-overlapping 100-bp windows and are shown as log_2_(counts□+□1). Chromosomes are resolved by parental species. **b,** Read-depth profiles across the assembled *D. simulans* centromeric contigs Cen2_3A_sim, Cen2_3B_sim, Cen4_sim and CenX_sim. **c,** Read-depth profiles across the assembled *D. melanogaster* centromeric contigs Cen2_mel, Cen3_mel, Cen4_mel and CenX_mel. Read depth in **b** and **c** is shown as log2(counts□+□1). Published repeat annotations, including satellite repeats, transposable elements and other repetitive-sequence classes, are shown below each centromeric contig.

### Asymmetric loss of D. melanogaster genomic material

Long-read nanopore sequencing of all four hybrid lines and both parental controls was performed. To maximize accuracy in repeat-rich and structurally variable regions, a new *D. melanogaster* reference genome was assembled from the Ras3 cell line using nanopore long-reads and Hi-C scaffolding **(Additional file 1: Fig. S1)**. The resulting assembly was evaluated by comparing major chromosome-arm lengths with dm6, assessing BUSCO completeness, examining genome-wide distributions of GC content, gene density, and repeat density, and inspecting Hi-C contact maps, which supported chromosome-scale structural integrity without evidence of major mis-assembly **(Additional file 1: Fig. S2a-d)**. For *D. simulans*, we used the w501 reference assembly available from NCBI RefSeq (GCF_016746395.2; Prin_Dsim_3.1), matching the paternal *D. simulans* genetic background used to generate the hybrid cell lines.

Read-depth analysis in 100 bp windows across all chromosomes revealed that parental controls are largely euploid, while all four hybrid lines display widespread aneuploidy with significant overrepresentation of *D. simulans* sequences **(Fig. 2a, Additional file 1: Fig. S3a)**. Most strikingly, all four lines have selectively lost the *D. melanogaster* allele of *Hmr* while retaining both alleles of the other characterized hybrid incompatibility genes (*Lhr*, *gfzf*, *Nup98-96*, and known HMR interactors) **(Additional file 1: Fig. S4).** Given that elevated HMR_mel_ causes cell division defects in hybrid backgrounds [4,5], this loss likely occurred during the early stages of cell line establishment and therefore provides powerful evidence for strong selective pressure against the HIC.

Beyond *Hmr_mel_* loss, each line independently underwent variable but directionally biased erosion of *D. melanogaster* sequences, predominantly affecting the X chromosome, chromosome 4, and chromosome 3R. HYC4 retains a near-complete complement of both genomes but shows selectively reduced *D. melanogaster* coverage of these chromosomes **(Fig. 2a, Additional file 1: Fig. S3b)**. HYC3 has lost most of the *D. melanogaster* X and chromosome 4, with read-depth evidence of a chimeric *D. melanogaster/D. simulans* fusion on chromosome 3R consistent with a trans-species recombination event **(Additional file 1: Fig. S5)**. HYC2 carries a truncated *D. melanogaster* X in stoichiometric balance with the *D. simulans* X, while HYC1 shows approximately half the *D. melanogaster* read depth across most autosomes, consistent with a partial *D. simulans* genome duplication **(Additional file 1: Fig. S3a)**. Mitochondrial DNA is exclusively of *D. melanogaster* origin in all lines, consistent with maternal inheritance **(Additional file 1: Fig. S6)**.

Centromeric regions show a parallel asymmetry: *D. simulans* centromeric islands are almost completely retained in all four hybrid lines, whereas *D. melanogaster* centromeric regions are partially eroded, most prominently for the X chromosome and chromosome 4 **(Fig. 2b,c)**. This erosion is consistent with the known enrichment of large, species-specific satellite blocks (such as the 359-bp repeat on the *D. melanogaster* X) that are absent in *D. simulans* and likely fail to be properly compacted by the hybrid nuclear environment [12,16,17].

### Dosage compensation of the hemizygous D. simulans X chromosome

Loss of the *D. melanogaster* X chromosome in HYC1 and HYC3 provides a natural experiment in aneuploidy management. Immunofluorescence using antibodies against MSL2, a core component of the Dosage Compensation Complex (DCC), revealed a distinct X-chromosomal territory in HYC1 and HYC3 but not in HYC4, which retains both X chromosomes **(Fig. 3a,b)**. This demonstrates that the *D. simulans* X chromosome, when rendered hemizygous, is competent to recruit the DCC and undergo dosage compensation. Clone HYC2, despite carrying a truncated *D. melanogaster* X, does not display a prominent MSL2 territory. We propose that the residual *D. melanogaster* X may retain enough DCC-docking sequences to prevent the *D. simulans* X from being scored as hemizygous, and additionally that the presence of chromosome 4 in HYC2 (substantially depleted in HYC1 and HYC3), which shares evolutionary history and sequence homology with the X chromosome [18], may titrate DCC components away from the *D. simulans* X.

**Fig. 3:**
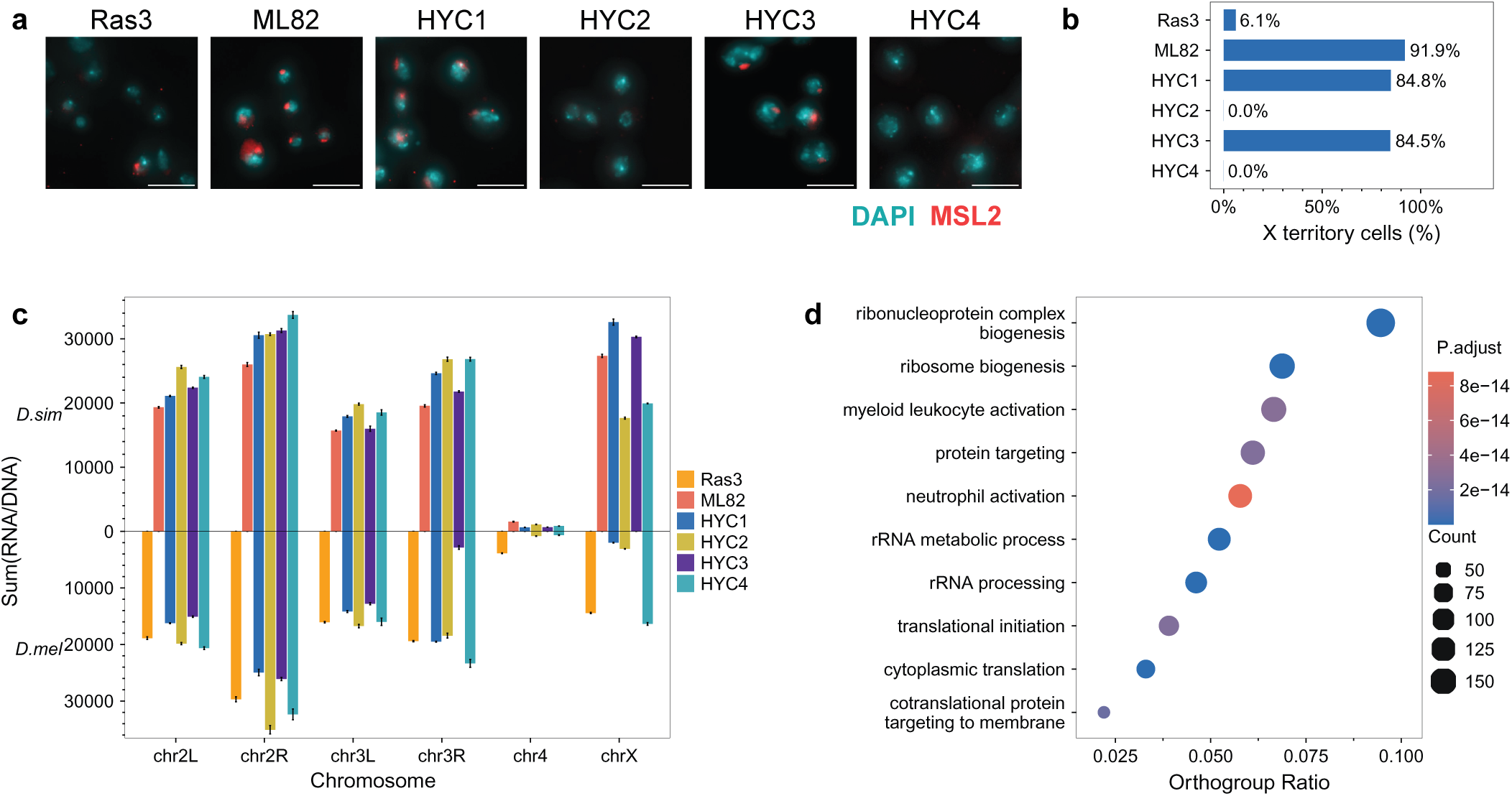
Dosage compensation and transcriptional responses in aneuploid hybrid cell lines. a, Representative immunofluorescence images of the parental control lines Ras3 (*Drosophila melanogaster*) and ML82 (*Drosophila simulans*) and the hybrid lines HYC1–HYC4 stained for MSL2 (red) and DNA (DAPI, cyan). Distinct MSL2-enriched chromosomal territories are visible in ML82, HYC1 and HYC3. Scale bars, 10 μm. **b,** Proportion of cells exhibiting a distinct MSL2-positive chromosomal territory. More than 200 cells were scored for each cell line. **c,** Chromosome-wide transcriptional output relative to genomic dosage. For each gene, sequencing-depth-normalized RNA abundance was divided by the corresponding normalized DNA abundance, and the resulting gene-level RNA/DNA values were summed for each parental chromosome. Values for *D. simulans*-and *D. melanogaster*-derived chromosomes are plotted in opposite directions from a common baseline to facilitate comparison of the two parental contributions, although all underlying RNA/DNA values are positive. Bars represent three biological RNA-sequencing replicates. **d,** Gene Ontology biological-process enrichment among orthogroups transcriptionally upregulated in the four hybrid cell lines analysed collectively relative to Ras3. Orthogroups with log2 fold change greater than 1 and Benjamini–Hochberg-adjusted *P□*<□0.05 were included in the enrichment analysis. Dot size indicates the number of input orthogroups assigned to each term, the horizontal axis shows the proportion of input orthogroups associated with the term, and colour indicates the adjusted *P* value.

To determine whether recruitment of the DCC was accompanied by increased transcription from the *D. simulans* X chromosome, we performed allele-resolved Oxford Nanopore RNA sequencing on three replicates of each hybrid and parental control cell line. RNA counts were normalized for library size, and normalized RNA abundance for each gene was divided by the corresponding sequencing-depth-normalized DNA abundance to estimate transcriptional output relative to genomic dosage. Gene-level RNA/DNA ratios were then summed across genes assigned to each chromosome.

The RNA/DNA signal of chrX_sim was approximately 1.5-fold higher in HYC1 and HYC3 than in HYC2 and HYC4 and was comparable to that in the *D. simulans* male control ML82 **(Fig. 3c)**. Together with the distinct MSL2-positive X-chromosomal territories in HYC1 and HYC3, this increase provides functional evidence for dosage compensation of the hemizygous chrX_sim in these two lines.

We next asked whether a comparable compensation mechanism operated on the aneuploid autosomes. Predicted protein sequences were analyzed with OrthoFinder to assign orthologous genes to orthogroups [19]. We then compared the *D. melanogaster* and *D. simulans* members of each orthogroup at the DNA, RNA, and RNA/DNA levels.

As expected, DNA abundance showed extensive chromosome-and clone-specific parental imbalance, which was closely mirrored at the RNA level **(Additional file 1: Fig. S7a-d)**. For example, chromosome 2L in HYC2 contained a higher expression of *D. simulans*-biased orthogroups, which was almost entirely based on differences in gene dosage as the bias was not detectable upon normalization of RNA abundance to the DNA dosage **(Additional file 1: Fig. S7e,f)**. A notable exception was the fourth chromosome which showed a strong bias towards expression of orthogroups from *D. simulans*. These results indicate that, with the exception of the fourth chromosome, parental expression biases on the autosomes were driven primarily by underlying copy-number differences rather than by compensatory allele-specific transcription.

We next investigated transcriptional responses shared across the independently derived hybrid lines. Relative to the *D. melanogaster* parental control Ras3, all four HYC lines showed elevated expression of orthogroups associated with ribosome biogenesis, translation and related ribosome functions **(Fig. 3d)**. We then compared the individual RNA outputs of the *D. simulans* and *D. melanogaster* members of each orthogroup within the hybrid lines. Gene Ontology analysis of orthogroups with higher transcriptional output from the *D. melanogaster* genes revealed recurrent enrichment of ribosome-related biological processes in HYC1–HYC4 **(Additional file 1: Fig. S8a–d)**. Thus, increased ribosome-related transcription was a shared feature of the hybrid cell lines and showed a recurrent bias towards the *D. melanogaster* alleles.

These transcriptional analyses reveal a generic response to extensive genomic imbalance observed in the hybrid lines. The chromosome-scale loss of the *D. melanogaster* X chromosome is compensated by the canonical dosage compensation of the retained *D. simulans* X, whereas a loss of individual genes or smaller regions of the *D. melanogaster* genome is largely compensated by a duplication of the corresponding *D.simulans* regions or not compensated at all. In general, the hybrid cells do not show major transcriptional aberrations when compared with each other or with the lines from pure species, which has also been observed in hybrid animals[20]. A notable exception is the recurrent elevation of ribosome-related genes, which identified ribosome production as a prominent and selectively regulated response to the hybrid cellular state.

### rDNA amplification and copy-number-dependent nucleolar dominance

As ribosomes also contain a substantial amount of RNA and as the number of rDNA genes in a cluster has been shown to be rather dynamic [21], we decided to also investigate the rDNA loci in the hybrid cell lines. To estimate parental rDNA copy number from genomic long-read data, we selected species-discriminating motifs from the 18S and 28S rDNA sequences of *D. melanogaster* and *D. simulans*. For normalization, we similarly designed species-specific motifs within seven pairs of X-linked single-copy BUSCO orthologs that remained detectable in the hybrid genomes, including the residual *D. melanogaster* X chromosome in HYC1 **(Additional file 1: Fig. S9)**. Motif occurrences were counted directly in the genomic reads using MotifScan, a tool we developed in this study. For each species and sample, the rDNA motif count was divided by the mean count of the corresponding seven X-linked BUSCO motifs, yielding an estimate of rDNA repeat abundance relative to one parental X-linked genomic complement.

Contrary to the global trend of *D. melanogaster* genome loss, all four hybrid lines show consistent and substantial enrichment of *D. melanogaster* rDNA copy number per X chromosome relative to the Ras3 parental control **(Fig. 4a, b)**. This increase is specific to rDNA repeat units and not observed in flanking single-copy regions, supporting locus-specific amplification. We propose that the intrinsically unstable, highly repetitive rDNA array is particularly susceptible to copy-number increase in the hybrid genomic background, analogous to rDNA magnification described in *Drosophila* [22,23].

**Fig. 4:**
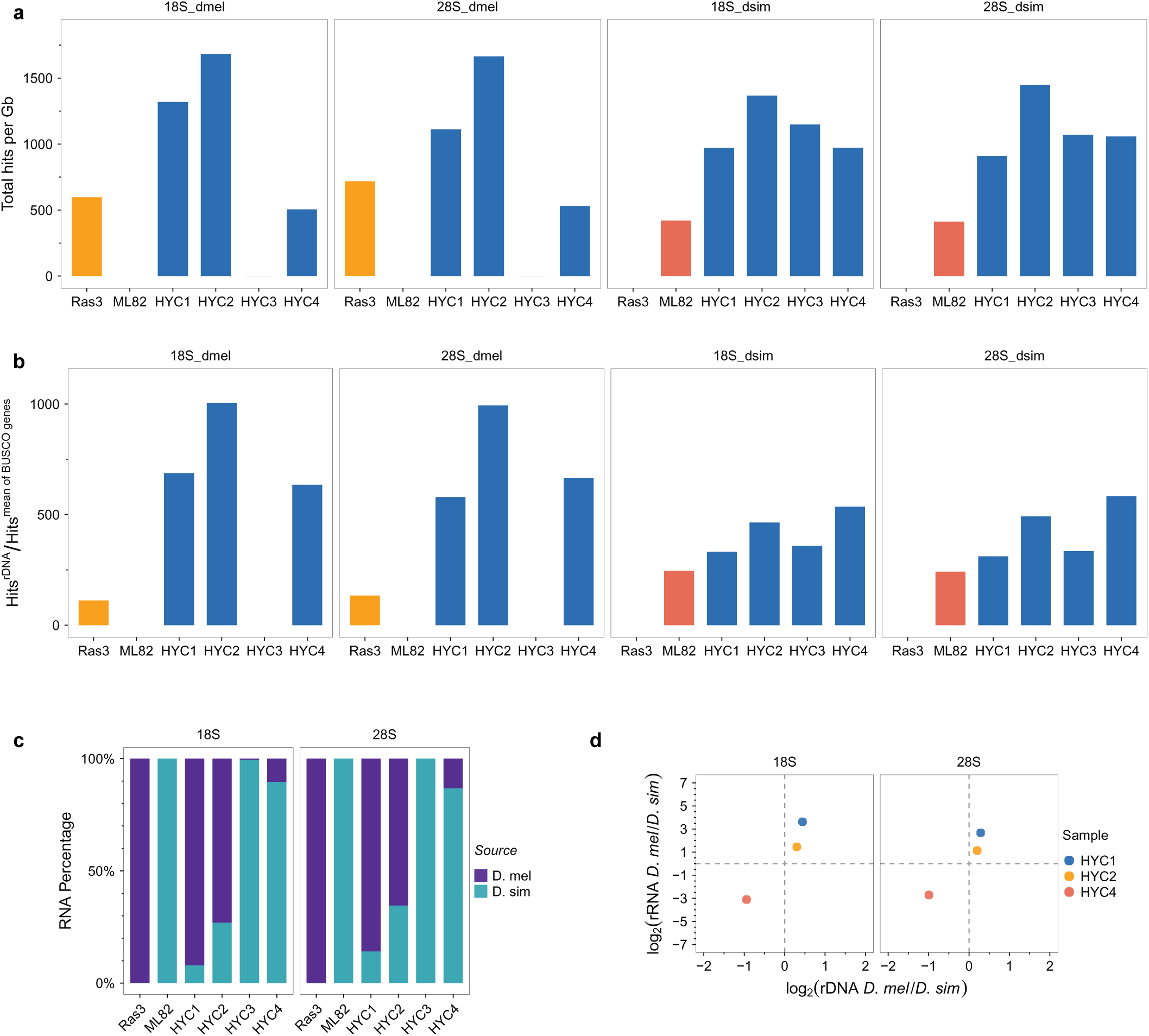
rDNA amplification and copy-number-dependent nucleolar dominance. a, Species-resolved abundance of 18S and 28S rDNA motifs in parental and hybrid cell lines. Motif counts were normalized to the total number of sequenced genomic bases and are expressed as motif occurrences per gigabase. **b,** Species-resolved estimates of 18S and 28S rDNA repeat abundance per parental X-linked genomic complement. rDNA motif counts were normalized to the mean signal from seven species-specific X-linked single-copy BUSCO orthologues. **c,** Mean species-of-origin proportions of mature 18S and 28S rRNA transcripts in the parental and hybrid cell lines. Values were averaged across biological RNA-sequencing replicates. **d,** Relationship between parental rDNA dosage and rRNA output in the hybrid cell lines. The horizontal axis shows log2(*D. melanogaster* rDNA/*D. simulans* rDNA), and the vertical axis shows the corresponding log2(*D. melanogaster* rRNA/*D. simulans* rRNA). Each point represents one hybrid cell line.

Analysis of rRNA transcript origin revealed that nucleolar dominance is not fixed to the D. melanogaster array as previously reported in whole-organism *D. melanogaster × D. simulans* hybrids [24,25]. Instead, Species-resolved 18S and 28S rRNA measurements revealed strong but line-specific transcriptional biases: *D. melanogaster*-derived rRNA predominated in HYC1 and HYC2, whereas *D. simulans*-derived rRNA predominated in HYC3 and HYC4 **(Fig. 4c)**. Importantly, the direction and magnitude of this transcriptional bias closely followed the relative genomic abundance of the parental rDNA copy number **(Fig. 4d)**. Thus, the observed rRNA asymmetry differs from a fixed, species-intrinsic form of nucleolar dominance. Instead, parental rRNA output appears to be largely dosage dependent, with the more abundant rDNA array contributing to the greater fraction of mature rRNA. These data indicate that the classical observation of *D. melanogaster* rDNA dominance in whole-animal hybrids likely reflects a dosage advantage rather than a fixed species-specific dominance signal, which is consistent with rDNA dosage manipulation switching nucleolar dominance in allopolyploid plants [26].

### Proteomic conflict as a novel driver of speciation

Next, we wanted to study how protein homeostasis is maintained when divergent genomes share a cellular environment. To minimize biases arising from unequal annotation depth between *D. melanogaster* and *D. simulans*, spectra were searched against a species-resolved protein database predicted from our long-read transcriptomes. Across all samples, we identified 106,591 peptide precursors with 7,394 peptides being only detected in hybrid lines (**Fig. 5a**). Of all detected peptides, 73.1% were shared between the two parental proteomes and could not be assigned unambiguously to either species, whereas 13.0% were specific to *D. melanogaster* and 13.9% to *D. simulans*. Because most peptides are thus shared between the two species, orthogroup-level protein abundance was quantified only from peptides whose sequence is present in both the *D. melanogaster* and the *D. simulans* member, which makes parental and hybrid measurements directly comparable and removes any dependence on species-specific peptide assignment. As has been reported before in many eukaryotes, the steady-state RNA abundance was only a weak predictor of protein abundance within individual cell lines. At the orthogroup level, transcript abundance explained only about 20% of the variation in protein abundance across the parental and hybrid lines **(Additional file 1: Fig. S10a)**. This limited correspondence is consistent with the substantial contribution of translational regulation and protein turnover to steady-state protein levels [27]

**Fig. 5:**
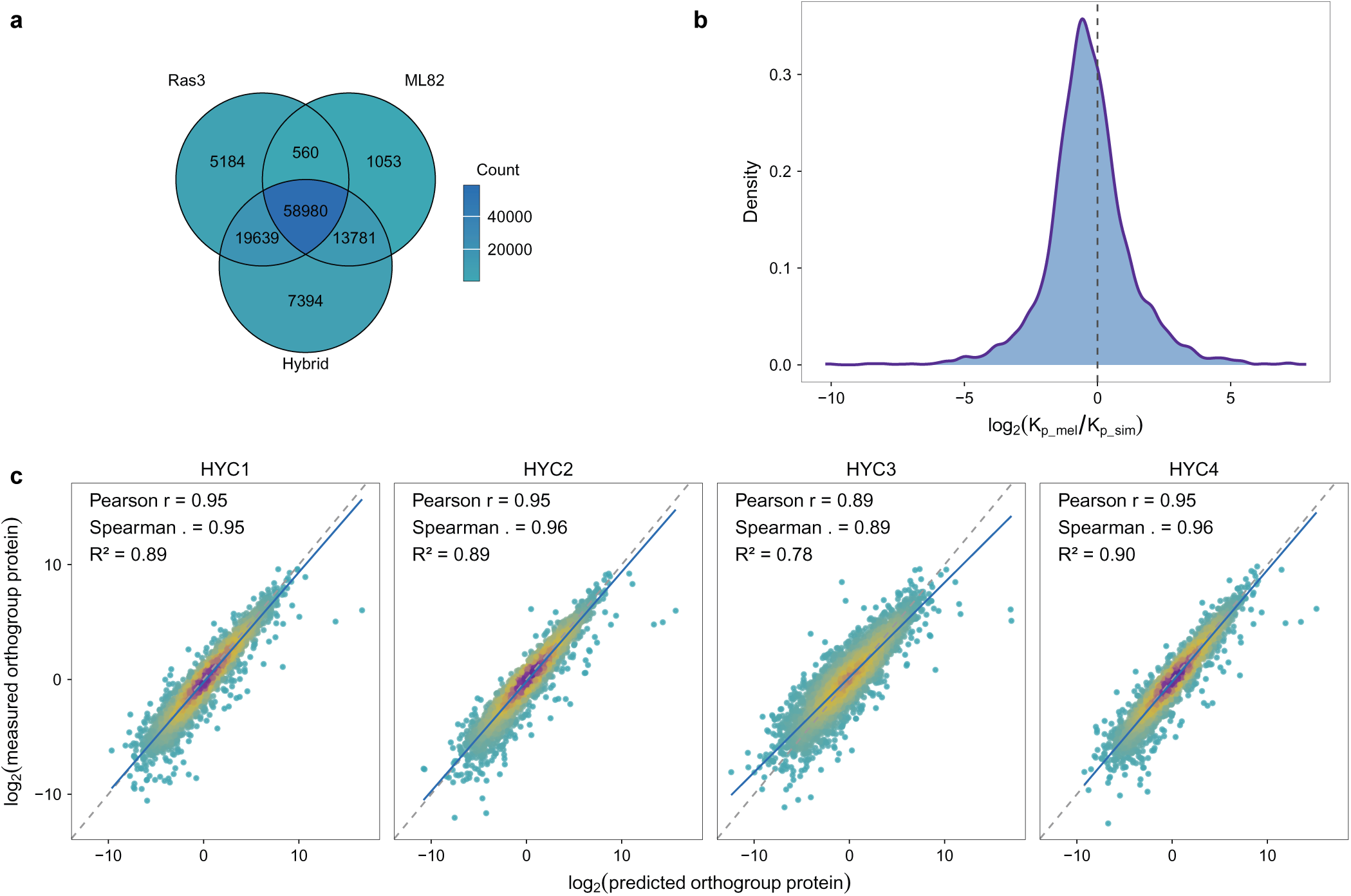
Parental protein-to-RNA relationships predict proteome output in hybrid cells. a, Overlap of distinct peptide sequences detected in Ras3, ML82 and the four hybrid cell lines considered as a single group. A peptide was classified as detected in a group when it was identified in at least one biological replicate from that group. Numbers indicate peptide sequences detected exclusively in each group or shared among groups. A total of 106,591 peptide precursors were identified across the complete DIA–MS dataset. **b,** Distribution of the relative post-transcriptional conversion factors between *D. melanogaster* and *D. simulans*. For each orthologous gene pair, K*_p_mel_* and K*_p_sim_* were defined as the protein-to-RNA ratios measured in Ras3 and ML82, respectively, and the distribution of log_2_(K*_p_mel_*/K*_p_sim_*) is shown. The dashed vertical line indicates a ratio of 1 (log = 0), corresponding to identical protein-to-RNA conversion between the two parental species. **c,** Predicted versus measured orthogroup-level protein abundance in HYC1–HYC4. For each gene g, species-specific protein-to-RNA conversion factors were estimated from the corresponding parental cell line, such that *K_p_mel_*= 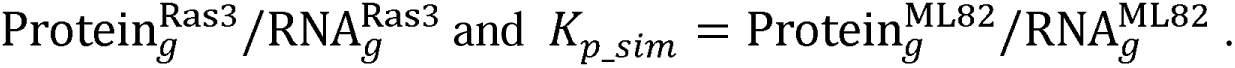 Predicted protein abundance for each hybrid orthogroup was calculated by applying the parental species-specific *K* values to the corresponding *D. melanogaster*-and *D. simulans*-derived RNA abundances in the hybrid and summing both parental contributions. Measured protein abundance represents the total orthogroup-level DIA–MS signal, for which species-specific protein contributions cannot generally be distinguished. Predicted and measured abundances are shown after log2 transformation. Dashed lines indicate the 1:1 relationship and solid lines indicate least-squares linear fits. Pearson r, Spearman *rho* and R² are indicated for each hybrid line.

To investigate whether the correlation between RNA and protein is hard wired into the sequence and independent of the cellular environment or whether it largely depends on the cellular translation and degradation machinery, we first calculated the steady-state translation-to-degradation ratio, termed K_p_ (where K_p_ = K_trans_/ K_deg_), for each protein in the pure species parental cell lines by dividing protein intensity to normalized RNA abundance (K_p_ = Protein/RNA). For most orthogroups, K_p_ values differed little between the two parental species, although the overall distribution showed a modest shift towards *D. simulans* **(Fig. 5b)**. Pairwise comparison of K_p_ values showed a relatively low correlation between the two parental lines, whereas the independently derived hybrid lines were generally more strongly correlated with each other **(Additional file 1: Fig. S10b).** Ranking orthogroups by the difference in parental *K_p_*further revealed a small subset of genes with pronounced divergence between *D. melanogaster* and *D. simulans* **(Additional file 1: Fig. S10c).**

We reasoned that if a protein’s K_p_ is primarily dictated by its intrinsic sequence rather than the overall trans-acting cellular composition, the total protein abundance of a given orthogroup () in the hybrid lines could be accurately predicted by integrating the allele-specific RNA transcript levels with their evolutionary parental K_p_ signatures:

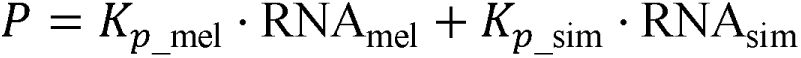

We observed a remarkably strong correlation between our calculated concentration values and the actual empirically measured protein abundances at the orthogroup level in the hybrids (**Fig. 5c**). This strong predictive power suggests that the K_p_ of a given protein is indeed intrinsically encoded in its sequence and evolves alongside it.

## Discussion

The four independently derived hybrid cell lines represent distinct molecular solutions compatible with sustained proliferation despite the coexistence of two incompatible genomes. Their endpoint genomes differed substantially, showing that hybrid cells do not converge on a single configuration. Nevertheless, the four lines share patterns of genome remodeling that reveal major molecular incompatibilities that cause postzygotic speciation.

By mapping which genomic regions were selectively purged across replicate lines, we can infer the relative strength of incompatibilities that accumulated during the divergence of *D. melanogaster* and *D. simulans*. In effect, the hybrid cell lines allow us to watch speciation run backwards at the cellular level. Each recurrent loss event represents a molecular fossil of an evolutionary conflict generated as independently evolving alleles and chromosomes became increasingly incompatible.

The convergence of independent lines on the same dominant constraint identifies *Hmr* as a major and rapidly co-evolved incompatibility in this system. HMR and LHR are central components of the incompatibility between *D. melanogaster* and *D. simulans*, and their mis-regulation disrupts chromatin organization and cell division in hybrid backgrounds. The uniform loss of *Hmr_mel_*, despite the distinct genomic architectures of the four lines and the retention of the other characterized incompatibility factors provides strong evidence that the different hybrid genomes repeatedly encountered the same major genetic constraint. Additional chromosome losses were more variable, but their shared bias toward the *D. melanogaster* chromosomes suggests that the *D. melanogaster* genome has accumulated more incompatibilities than the genome of *D.simulans* or that is is easier to compensate the loss.

The fact that genome erosion of the *D. melanogaster* is not happening at random allows us to predict the strongest genomic incompatibilities between the two genomes. The most frequent loss of genetic material occurs on the D. *melanogaster* X-chromosome, which in two lines results in an activation of the dosage compensation complex and subsequently to an active dosage compensation of the hemizygous *D. simulans* X. Interestingly, the rDNA locus that is located on the X-chromosome experiences a magnification on the remaining *D. melanogaster* X-chromosome as well as on the *D. simulans* X-chromosome. This magnification is only observed in hybrid cells but not in cell lines derived from pure species, suggesting that it is caused by their hybrid nature rather than the establishment of stable tissue culture lines. Similar to what has been reported in interspecies hybrid flies [25,28], we observe a strong nucleolar dominance. However, in contrast to what is seen in flies, we observe dominance of the locus that contains more rDNA copies rather than the one derived from *D. melanogaster*. This result suggests that nucleolar dominance might be determined by the copy number rather than the species origin.

Besides the genomic alterations our analyses also revealed the differences in the RNA to protein coefficient K_p_ as a novel and yet undescribed source of molecular conflict in hybrid species. This is in agreement with the puzzling previous observation of whole-transcriptome analyses of *D. melanogaster D. simulans* hybrids which showed surprisingly modest global RNA expression differences compared to the parental species [29] suggesting that any potential incompatibility is rather due to differences in posttranscriptional regulation than changes to a mis-regulation of transcription. In fact, that fits well to our previous observation that hybrid incompatibility is caused an aberrantly high abundance of the *D. melanogaster* HMR protein, which disrupts chromatin organization[4]. The lethality driven by HMR is therefore not a symptom of global transcriptional collapse, but rather a localized failure caused by a vastly different, species-specific intrinsic RNA to protein coefficient of *Hmr* expression. In hybrids toxic protein levels of an HMR/LHR complex are produced that are normally prevented by a robust buffering system in pure species. Thus, the forced coexistence of these genomes leaves the cells with only one viable solution to bypass this stoichiometric bottleneck: the physical elimination of the *D. melanogaster Hmr* allele.

## Conclusions

Our studies of hybrid cell lines show that evolutionary incompatibilities can arise across all steps of the central dogma from chromosome organization to transcription and translation efficiency. By interrogating hybrid survival at a molecular resolution, our findings demonstrate that speciation barriers are not restricted to regulatory mismatches in DNA or RNA but can also manifest as differences within the proteome. The HYC1–HYC4 cell lines provide an experimental platform to decipher these rules. While these lines represent viable end-states rather than intermediate stages, their convergent rearrangements reveal that hybrid genome stabilization is neither an unconstrained collapse nor a single deterministic trajectory. Instead, hybrid genomes resolve along defined trajectories shaped by physical gene dosage, translational capacity, and the necessity of preserving protein homeostasis

## Methods

### Fly strains and cell line generation

Hybrid cell lines were generated following a modified Simcox protocol [13,14]. Virgin female *Drosophila melanogaster* carrying Act5C-GAL4 in a yw background (genotype: y1 w*; P{Act5C-GAL4}25FO1/CyO, y+), were crossed to male *Drosophila simulans* carrying UAS-RasV^12^ in an w501 background (genotype, UAS-Ras^V12^/UAS-Ras^V12^). Overnight embryos were collected, dechorionated, allowed to cellularise, and homogenized before being transfered into T25 culture flasks containing Schneider’s *Drosophila* medium supplemented with 10% heat-inactivated fetal bovine serum (FBS; Capricorn#FBS-12A) and 100 units/ml penicillin-streptomycin (Sigma#P0781). Cultures were maintained at 23°C. Confluent cultures were passaged following treatment with 0.2% trypsin. Immortalized cell lines were considered established once cultures grew continuously and could be passaged reproducibly; this required approximately 3-7 months. Four independently derived hybrid cell lines were established and designated HYC1, HYC2, HYC3, and HYC4. The *D. melanogaster* parental control line Ras3 was generated from *D. melanogaster* Act5C-GAL4 × UAS-Ras^V12^ crosses as described previously[15], was obtained from the laboratory of Yuri B. Schwartz. The *D. simulans* parental control line ML82-19a was obtained from the *Drosophila* Genomics Resource Center (DGRC#CVCL_Z802).

### DNA-FISH (DNA Fluorescence in Situ Hybridization)

DNA Probes used in study are provided by Madhav Jagannathan and have been previously described [16,30] *D. mel* specific probe (AATAACATAG), *D. sim* specific probe (GAACAGAACATGTTCGAACAGAACATGTTCGAACA), and *D. mel/D. sim* common probe (AATAT)_6_.

DNA-FISH were performed as described previously [16,31]. Briefly, cells were cultured on coated cover slips for one day. The medium was removed, and the cells were fixed with 4% formaldehyde in PBS for 10 min at room temperature. After rinsing with PBS, the cells were permeabilized with ice-cold PBS containing 0.4% Triton X-100 for 5 min on ice. The cells were immediately rinsed and washed twice with PBS and then blocked with 5% normal goat serum (NGS; Jackson ImmunoResearch, catalog no. 005-000-121) in PBS for 30 min. After rinsing with PBS, the cells were incubated overnight at 4°C with an anti-lamin antibody diluted 1:500 in PBS containing 5% NGS. The cells were washed once for 5 min with PBS containing 0.1% Triton X-100, followed by three washes with PBS. They were then incubated for 1 hour at room temperature with goat anti-mouse Alexa Fluor 647 secondary antibody (1:1000; Invitrogen, catalog no. A-21236) for samples hybridized with the *D. melanogaster/D. simulans* common probe or the *D. simulans*–specific probe, or with donkey anti-mouse Alexa Fluor 488 secondary antibody (1:500; Jackson ImmunoResearch, catalog no. 715-545-140) for samples hybridized with the *D. melanogaster*–specific probe. After three washes with PBS, the cells were post-fixed with 3.7% paraformaldehyde in PBS for 10 min at room temperature in the dark. The cells were rinsed three times with PBS and incubated with 0.1 μg/μl RNase A in PBS for 1 hour at 37°C. After three additional rinses with PBS, cellular DNA was denatured by incubation in 1.9 M HCl for 30 min at room temperature. The cells were rinsed three times with PBS and hybridized overnight at 37°C in a dark, humidified chamber with the corresponding probe in hybridization buffer containing 2× SSC (0.3 M NaCl and 0.03 M sodium citrate), 500 nM probe, and 60% formamide. Coverslips were flooded with 2× SSC and washed three times with 2× SSC. DNA was counterstained with 0.2 ng/ml DAPI for 5 min, followed by rinsing with 2× SSC. Coverslips were mounted in mounting medium and sealed with nail polish.

Confocal microscopy was performed at the Core Bioimaging Facility of the Biomedical Center using an inverted Leica SP8 confocal microscope equipped with 405-, 488-, 552-, and 638-nm excitation lasers. Images were acquired using a 100×, 1.4–numerical aperture objective at a pixel size of 80 nm. DAPI, Cy3, Alexa Fluor 488, and Alexa Fluor 647 fluorescence was detected using the corresponding acquisition channels. Sequential scanning was used to minimize spectral bleed-through. Cy3, Alexa Fluor 488, and Alexa Fluor 647 signals were detected using hybrid detectors, whereas DAPI fluorescence was detected using a conventional photomultiplier tube. The resulting.lif files were processed using Fiji v1.54f [32]. More than 50 cells were imaged and analyzed for quantification of probe foci.

### Nanopore long-read genomic sequencing

One million cells were collected by centrifugation, washed twice with PBS and stored at-80°C until use. Genomic DNA was isolated from the cell pellet with Qiagen Blood & Cell Culture DNA MiniKit (Qiagen#13323) according to the manufacturer’s protocol. A total of 1 ug of DNA was further purified with 0.6 volumes of AMPure XP beads (Beckman-Coulter#A63881) and used for preparation of nanopore sequencing libraries with the Native Barcode Ligation kit (Oxford Nanopore Technologies#NBD114.96) according to the manufacturer’s protocol. Barcoded libraries were quantified, pooled at equimolar ratio and sequenced on a PromethION P24 instrument using FLO-PRO114M flow cells for 72 h. Fastq files were generated by MINKNOW real-time base calling with the superaccurate model.

### Assembly of Ras3 genome

The genome assembly workflow is summarized in **Additional file 1: Fig. S1**. NanoQC v0.10.0 [33] was used to assess the quality of the raw Nanopore sequencing reads generated from the Ras3 cell line. Raw fastq.gz files were filtered with chopper v0.8.0 to retain reads with a mean sequencing quality score of at least 20 and a minimum length of 5 kb [33]. The filtered reads were assembled *de novo* into contigs using flye v2.9.3-b1797[34]. Consensus correction was performed using medaka v1.11.3 (https://github.com/nanoporetech/medaka, from Nanopore Tech). The consensus contigs were polished with racon v1.5.0[35] for three rounds. The *Drosophila melanogaster* 6.56 genome from FlyBase[36] was used as a reference in Ragtag v2.1.0[37] to guide the scaffolding of contigs. TGS-GapCloser v1.2.1[38] was used to close gaps in the draft assembly. Hi-C data of Ras3 can be accessed from GEO datasets with the accession number GSE198763[15]. Hi-C data was mapped to the draft assembly with bwa-mem2 v2.2.1 [39] to rearrange the genome with Juicer v1.6[40] and 3D-DNA v180114 [41]. Rearranged chromosomes were manually inspected and corrected with JuiceBox v2.17.00[40]. Two iterative rounds of 3D-DNA post-review scaffolding and manual correction in Juicebox were performed to generate the final chromosome-scale assembly. The assembled Ras3 genome is provided as Data S1.

Primary genome annotation was performed with liftoff v1.6.3[42] using the annotation information of *Drosophila melanogaster reference genome release 6.56* from FlyBase [36] as a reference. Primary repeat annotation was performed with RepeatMasker v4.1.8 (https://www.repeatmasker.org/) using the Dfam database as reference.

The completeness of the assembled Ras3 genome was evaluated using Benchmarking Universal Single-Copy Orthologs (BUSCO) v5.6.1[43]. The analysis was performed against the *eukaryota_odb10* lineage dataset in genomic mode. To assess the structural integrity and contiguity of the assembled genome, Hi-C data mentioned above were utilized to visualize the three-dimensional (3D) chromatin organization. Hi-C data was trimmed with Trim-Galore v0.6.10[44] and then mapped to Ras3 genome with bwa-mem2 v2.2.1[39]. Mapping result was processed with pairtools v1.1.0[45] with mapping quality above 40. Contact matrix was generated as.cool and.mcool files with cooltools v0.7.1 [46]. Visualization was performed with HiContacts v1.4.0 [47] and custom scripts.

### Analysis of long-read genomic sequencing data

Nanopore long reads were first assessed for quality using NanoQC[33]. Reads were subsequently filtered using Chopper v0.8.0 [33] to retain sequences at least 2 kb in length with a mean sequencing quality score of at least 20. An additional 50 bp was trimmed from each end of every read. Filtered reads were aligned to reference genomes using minimap2 v2.26-r1175[48] with parameters *-ax map-ont*.

For genome-wide coverage analysis, a combined reference genome was constructed by merging the *D. melanogaster* assembly (Ras3, generated in this study) with the *D. simulans* reference genome (Dsim3.1, NCBI accession: GCF_016746395.2). For centromeric coverage analysis, a combined reference was generated using centromeric contigs of *D. melanogaster* or *D. simulans* from Cécile Courret et al[17]. For mitochondria coverage analysis, a combined mitochondrial reference was constructed by combining mtDNA from Ras3 and mtDNA (NCBI accession: CM028646.2) from Dsim3.1.

Aligned reads were processed using samtools v1.21-17-g9e91173[49] to retain high-confidence mappings. For genome-wide and mitochondria analyses, reads with mapping quality ≥ 30 were retained, whereas for centromeric regions, a relaxed threshold of mapping quality ≥ 1 was applied due to the repetitive nature of these sequences. Genome-wide coverage profiles were computed by generating bin-based coverage matrices using deepTools v3.5.6[50] with specific bin size. In order to estimate the ploidy, we calculated the genome-wide bin-based coverage matrices using deepTools v3.5.6 [50] with a bin size of 100 bp.

Fusion event in HYC3 cell line was identified through manual inspection of read alignments in Integrative Genomics Viewer (IGV) v2.19.7[51]. Reads spanning breakpoint at 3R_mel were extracted and split at alignment discontinuities, and segments were independently realigned to the combined reference genome. The fusion was supported by mapping split segments from the same read to *D. melanogaster* 3R and *D. simulans* 3R. The breakpoint was supported by at least nine independent long reads.

### Species-resolved long-read RNA sequencing

Total RNA was extracted from one million cells using the Zymo Quick-RNA Miniprep Kit (ZYMO#R1054) according to the manufacturer’s protocol. 500 ng of total RNA was used for preparation of full-length cDNA libraries with the cDNA PCR barcoding kit (Oxford Nanopore Technologies# PCB114.24) according to the manufacturer’s protocol. Briefly, mRNA was reverse transcribed by polyA-specific priming, full-length cDNA was selected by a template-switching mechanism and PCR amplification of double-adapted molecules. Barcoded libraries were quantified, pooled at equimolar ratio and sequenced on a PromethION P24 with FLO-PRO114M flowcells for a runtime of 72 h. Fastq files were generated by MINKNOW real-time base calling with the superaccurate model.

Reads were assessed with NanoQC [33] to inspect read length distribution, per-read quality scores and overall sequencing performance. Reads were then filtered with chopper to retain high-quality reads with a mean sequencing quality score ≥20. Filtered reads were mapped to a combined species reference genome consisting of the Ras3 assembly generated in this study and the Dsim3.1 reference genome using minimap2 v2.26-r1175[48] with splice-aware parameters *-ax splice*. The resulting alignments were sorted and indexed with samtools v1.21-17-g9e91173 [49]. To minimize ambiguity between species-derived transcripts, only high-confidence alignments with mapping quality ≥30 were retained for downstream analyses. The filtered BAM files were used as input for Isoquant v3.7.1[52] to reconstruct transcript models and quantify gene-level expression. The reference annotation supplied to IsoQuant was generated by combining the primary gene annotation of the Ras3 assembly with the gene annotation of Dsim3.1.

To generate a unified protein-coding gene set for orthology and functional annotation analyses, transcript sequences were extracted from the IsoQuant annotation, and coding sequences were predicted and translated into protein sequences using TransDecoder2 v1.0.6[53]. Protein sequences were then assigned to their species of origin according to whether the corresponding genes mapped to the Ras3-derived *D. mel* reference or to the Dsim3.1-derived *D. sim* reference. Orthogroups between the two species-derived protein sets were inferred using OrthoFinder v3.1.0 [19]. For each orthogroup, IsoQuant-derived gene expression values were summed separately for *D. mel* and *D. sim* members to generate species-resolved orthogroup expression estimates.

For each orthogroup, the *D. mel* expression fraction was calculated as *D. mel* expression divided by the sum of *D. mel* and *D. sim* expression. Orthogroups were classified as *D. mel*-biased, *D. sim*-biased or unbiased using thresholds of >0.7, <0.3 or intermediate values, respectively.

Protein sequences were annotated using eggNOG-mapper v2.1.13[54]. GO annotations for each orthogroup were defined as the union of GO terms assigned to all proteins within that orthogroup. GO enrichment analyses were performed using clusterProfiler v4.18.4[55], with all expressed orthogroups as the background universe. For allele-specific differential analyses, RNA/DNA ratios of the *D. melanogaster* and *D. simulans* members of each orthogroup were compared using limma v3.66.0[56], P values were adjusted using the Benjamini–Hochberg procedure. Orthogroups with log2FC (*D. melanogaster/D. simulans*) > 1 and adjusted P < 0.05 were classified as D. melanogaster-biased and subjected to Gene Ontology enrichment analysis. GO terms with Benjamini–Hochberg-adjusted P values below 0.05 were considered significantly enriched.

### MSL2 staining

Immunofluorescence staining was performed on Ras3, ML82-19a, and HYC1–HYC4 cells grown on glass coverslips. Briefly, one million cells were pipetted onto 12-mm glass coverslips and allowed to settle for 12 hours in a 12-well plate. Coverslips were then transferred to a clean 12-well plate and washed with PBS for 10 min at room temperature. Cells were fixed for 10 min at room temperature in PBS containing 3.7% methanol-free formaldehyde, rinsed three times with PBS, and washed twice for 5 min in PBS. Cells were permeabilized for 6 min on ice with ice-cold PBS containing 0.25% Triton X-100, rinsed three times with PBS, and washed once for 5 min in PBS. Coverslips were blocked for 1 h at room temperature in PBS containing 5% NDS (Normal Donkey Serum, Jackson Immuno Research#017-000-121). Coverslips were incubated overnight at 4°C with rabbit anti-MSL2 primary antibody (provided by Peter Becker’s Lab) diluted 1:800 in PBS containing 5% NDS. Before application, antibody dilutions were centrifuged for 5 min at 15,000 rpm at 4°C. Coverslips were washed for 10 min with PBS containing 0.1% Triton X-100, followed by an additional PBS wash. Coverslips were then incubated for 1 h at room temperature with Alexa Fluor 647-conjugated donkey anti-rabbit secondary antibody (Jackson Immuno Research#711-605-152) diluted 1:200 in PBS containing 5% NDS. From this step onward, samples were protected from light. Coverslips were washed for 5– 10 min with PBS containing 0.1% Triton X-100, followed by an additional PBS wash. DNA was counterstained with DAPI diluted 1:500 in PBS for 5 min, followed by two 5-min washes in PBS. Finally, coverslips were mounted using 4 µl mounting medium per coverslip and sealed with nail polish. Slides were stored at 4°C protected from light until imaging.

Widefield image stacks were recorded with a HC PL APO 100x/1.40 oil objective on a motorized Leica DMi8 inverted microscope with Spectra X LED light source from Lumencor. Camera: Hamamatsu ORCA-Flash 4.0; Image pixel size: 65 nm. Plane spacing: 300 nm. Fluorescence images were recorded with a quadband filter, selective LED illumination and an emission filter wheel to avoid channel blead-through. Effective excitation and emission bandwidth (Ex/Em): DAPI: 379-401 nm/ 450-490 nm; Alexa 647: 625-655 nm/ 670-770 nm. Software: LAS X 3.7.6.25997. Image stacks were directly maximum projected in the LAS X software and then saved in the vendor’s original format as.lif. Obtained.lif files were processed with Fiji v1.54f[32]. For quantification of MSL2 staining patterns, more than 200 cells were imaged and analyzed.

### Species-resolved rDNA copy number and rRNA origin analysis

To quantify rDNA copy number and rRNA origin in a species-resolved manner, we used diagnostic SNPs that distinguish *D. melanogaster* and *D. simulans* rDNA sequences. The 18S and 28S rDNA sequences of the two species were aligned, and species-informative SNPs were identified within the mature rRNA regions. Diagnostic sequence motifs containing these SNPs were then used to distinguish *D. melanogaster*– and *D. simulans*–derived rDNA and rRNA molecules.

For rDNA copy-number estimation, high-quality nanopore genomic reads were scanned for diagnostic 18S and 28S rDNA motifs using MotifScan v0.1.7, a custom high-throughput motif-scanning tool developed for rapid detection of user-defined diagnostic sequence motifs in long-read sequencing data (https://github.com/jiehua1995/MotifScan). Species-specific rDNA hits were counted separately for 18S and 28S. To normalize these counts, we selected seven X-linked single-copy BUSCO genes whose *D. melanogaster* and *D. simulans* alleles could also be distinguished by species-informative SNPs. Reads containing diagnostic motifs from these BUSCO genes were counted using the same procedure as for rDNA. Diagnostic motif hits within these BUSCO genes were counted using the same procedure as for rDNA. For each species and sample, the mean hit count across the seven X-linked BUSCO genes was used as a single-copy normalization factor.

This calculation was performed separately for 18S and 28S rDNA, and independently for *D. melanogaster* and *D. simulans* alleles. Because the normalization loci are X-linked and single-copy, the resulting values estimate rDNA abundance relative to one species-specific X-linked genomic equivalent.

This normalization strategy is conservative in the context of the hybrid cell lines. The *D. melanogaster* orthologues of the seven X-linked BUSCO genes are retained in HYC1, HYC2 and HYC4 despite substantial loss of *D. melanogaster* X-chromosomal sequence in HYC2. As a result, *D. melanogaster* rDNA enrichment is measured relative to the residual retained *D. melanogaster* X-linked genomic complement rather than against total genome-wide coverage, preventing inflation of rDNA enrichment estimates caused by broad *D. melanogaster* genomic erosion.

Although the cDNA libraries were generated using poly(A)-specific priming, substantial numbers of reads derived from mature 18S and 28S rRNAs remained detectable in all libraries. These reads were used exclusively to determine the relative species of origin of recovered rRNA molecules and not to compare absolute rRNA abundance among libraries. High-quality nanopore reads were scanned using MotifScan v0.1.7 for the same species-discriminating 18S and 28S motifs used for rDNA analysis. Motif hits were counted separately for *D. melanogaster* and *D. simulans* in each biological replicate. For each rRNA species, the *D. melanogaster* and *D. simulans* proportions were calculated relative to the total number of species-assigned motif hits. Measurements were performed independently for 18S and 28S rRNA.

### Proteomics sample preparation, DIA–MS analysis and data processing

Approximately 1 x 10^6^ cells were collected for each biological replicate, with three biological replicates prepared for Ras3, ML82 and each of the four hybrid cell lines. Protein extraction, reduction, alkylation, enzymatic digestion and peptide preparation were performed using the PreOmics iST 8x kit (PreOmics, P.O.00001) according to the manufacturer’s instructions.

For LC–MS/MS purposes, desalted peptides were injected in an Ultimate 3000 RSLCnano system (Thermo), separated in a 25 cm analytical column (75 µm ID, 1.6 µm C18, Aurora-IonOpticks) with a 50-minute gradient from 2 to 35% acetonitrile in 0.1% formic acid. The effluent from the HPLC was directly electrosprayed into an Orbitrap Exploris 480 (Thermo) operated in data-independent mode to automatically switch between full scan MS and MS/MS acquisition of 30 windows of 20 m/z each, covering the range of 380-980 m/z with 1 m/z overlap. Survey full scan MS spectra (from m/z 380 to 980) were acquired with resolution R = 120,000 at m/z 400 (AGC target of 3 × 10^6^) and max IT of 100ms. DIA scans were acquired with R = 30000 (AGC target of 3 × 10^8^), max IT set to “Auto” and normalized collision energy set at 30%. Typical mass spectrometric conditions were as follows: spray voltage, 1.5 kV; heated capillary temperature, 275°C.

To minimize biases resulting from the unequal completeness of the public *Drosophila melanogaster* and *Drosophila simulans* protein annotations, spectra were searched against a species-resolved protein database generated from the long-read transcriptomes produced in this study. Transcripts reconstructed with IsoQuant were translated using TransDecoder, and the resulting *D. melanogaster* and *D. simulans* protein sequences were combined into a single FASTA database while retaining species-specific transcript and gene identifiers. Raw DIA files were analysed using DIA-NN v2.2.0 Academia in library-free mode with in silico spectral-library prediction. Searches assumed tryptic cleavage after lysine or arginine and allowed one missed cleavage. Peptides of 7–30 amino acids and precursor charge states of 1–4 were considered. Carbamidomethylation of cysteine was specified as a fixed modification and oxidation of methionine as a variable modification. Precursor and fragment mass ranges were 300–1,800 *m*/*z* and 200–1,800 *m*/z, respectively. Identifications were filtered at a 1% false-discovery rate. Protein inference was disabled to retain peptide-level information and avoid assigning peptides shared between parental orthologues to a single inferred protein group.

Peptide-level quantities were processed in R. Peptides were classified as *D. melanogaster*-specific, *D. simulans*-specific or shared according to the species identities of all protein sequences to which they mapped. Peptides mapping exclusively to proteins from one species were classified as species-specific, whereas peptides mapping to proteins from both species were classified as shared. Peptide detection across Ras3, ML82 and the combined hybrid-cell group was summarized from distinct peptide precursors. Peptide-level intensities were also used for principal-component analysis after log2 transformation and filtering of peptides with insufficient quantitative observations across samples.

For orthogroup-level analyses, protein and gene identifiers associated with each peptide were linked to the orthogroups inferred by OrthoFinder from the transcriptome-derived protein sequences. Peptides mapping to more than one orthogroup were excluded for each of those orthogroups. To avoid any dependence on species-specific peptide assignment and to make the parental and hybrid measurements directly comparable, only strict 1:1 orthogroups (exactly one *D. melanogaster* and one *D. simulans* member) were used, and orthogroup protein abundance was quantified exclusively from common peptides, defined as peptides whose exact amino-acid sequence is present in both the *D. melanogaster* and the *D. simulans* member protein of that orthogroup. Within each replicate, the intensities of all common peptides assigned to an orthogroup were summed; the resulting value was divided by the median positive orthogroup value of the same replicate to correct for loading differences, and the three replicates were averaged. Parental and hybrid libraries were normalized independently by the same procedure. A common peptide was considered detected in a parental line when it was positive in at least one replicate. An orthogroup was retained only if it was supported by at least two distinct common peptide sequences and every retained common peptide was detected in both parents. Missing values were treated as not detected and contributed nothing to the sums; values were not imputed. Orthogroup abundances were log2-transformed before statistical analysis.

For integration of the proteomic and transcriptomic datasets, RNA abundance was likewise summarized at the orthogroup level by combining the expression of genes assigned to the same orthogroup. Orthogroups with quantitative measurements in both datasets were retained for comparison. RNA and protein abundances were log2-transformed before statistical analysis. Within each cell line, the relationship between RNA and protein abundance was assessed by linear regression across matched orthogroups, and the coefficient of determination (R^2^) was used to quantify the proportion of variation in protein abundance associated with RNA abundance.

For each orthogroup, a steady-state protein-to-RNA ratio, was calculated in the Ras3 and ML82 parental cell lines as K_p_ = *P/R*, where *P* and *R* denote the orthogroup-level protein and RNA abundances, respectively.

To test whether parental protein-to-RNA relationships could predict protein abundance in the hybrid cell lines, species-resolved RNA abundances in each hybrid were combined with the corresponding parental *Kp_mel_ or Kp_sim_* values. For each orthogroup, the predicted total protein abundance was calculated as

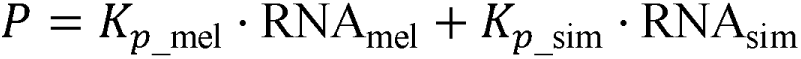

The predicted contributions of the two species were summed because most detected peptides could not be unambiguously assigned to one parental orthologue and therefore provided a combined orthogroup-level protein measurement. Predicted total protein abundances were compared with the empirically measured orthogroup-level protein abundances in HYC1–HYC4 using Pearson correlation, Spearman rank correlation and linear regression.

## Declarations

### Ethics approval and consent to participate

Not applicable.

### Consent for publication

Not applicable.

### Availability of data and materials

Long-read genomic and transcriptomic sequencing data have been deposited in the NCBI Sequence Read Archive under BioProject accession PRJNA1216052. The associated records are currently under embargo and will be made publicly available upon acceptance of the manuscript. The mass spectrometry proteomics data have been deposited with the ProteomeXchange Consortium through the PRIDE [57] partner repository under dataset identifier PXD081377.

Data S1–S5, comprising the Ras3 chromosome-scale genome assembly, the combined *Drosophila melanogaster*–*Drosophila simulans* reference genome and annotation, IsoQuant-derived coding sequences and predicted protein sequences, have been deposited in Zenodo under the reserved DOI 10.5281/zenodo.20760248. The Zenodo record is available to editors and reviewers through a private review link and will be made publicly available upon acceptance of the manuscript. The hybrid cell lines are available upon request to the corresponding author. All other data supporting the findings of this study are available in the Article, its additional files.

### Code availability

MotifScan, the sequence motif-scanning software developed in this study, is publicly available at: https://github.com/jiehua1995/MotifScan. Custom scripts and Quarto notebooks used for genomic, transcriptomic, proteomic and integrative analyses, and for reproducing the figures in this study, are publicly available at: https://github.com/jiehua1995/Hybrid_Cells

## Acknowledgments

We thank Madhav Jagannathan and Yukiko M. Yamashita for providing DNA-FISH probes; Peter Becker for anti-MSL2 antibody; Marisa Müller for immunofluorescence protocol assistance. We acknowledge the Laboratory for Functional Genome Analysis (LAFUGA), Gene Center Munich, LMU Munich, for sequencing support; the Core Facility Bioimaging, Biomedical Center, LMU Munich, for microscopy and imaging support; and the Zentrallabor für Proteinanalytik (ZfP; Protein Analysis Unit), LMU Munich, for mass spectrometry-based proteomic analysis. We thank all members of the Imhof laboratory for scientific discussions.

## Author contributions

J.H. and A.I. conceived the study. J.H., A.L. and T.G.K. developed the methodology. J.H., A.L., T.G.K., A.W.T. and R.C. performed the experiments. J.C. generated the *D. simulans* strain used in this study. I.F. performed the mass spectrometry measurements. J.H. performed the formal analyses, curated the data and generated the visualizations, with input from A.I. N.P., Y.B.S. and A.I. provided resources. Y.B.S. and A.I. acquired funding. J.H. and A.I. administered the project. A.I. supervised the study. J.H. and A.I. wrote the original draft. All authors reviewed and edited the manuscript.

## Supplementary Figure Legends

**Fig. S1. Genome assembly pipeline for the Ras3 D. melanogaster cell line.** Raw Nanopore reads were assessed using NanoQC and NanoPlot, adapter-trimmed and filtered with Chopper, and assembled de novo with Flye. The initial contigs were polished with Medaka and three rounds of Racon. Redundant haplotigs were identified using RagTag using the *D. melanogaster* dm6 reference genome to guide assembly curation. Remaining assembly gaps were closed with GapCloser. Hi-C reads were processed with Juicer and 3D-DNA and used to scaffold the curated contigs, followed by manual refinement with Juicebox to generate the final Ras3 genome assembly. Assembly completeness was evaluated using BUSCO. Gene annotation was transferred with Liftoff, and repetitive sequences were annotated with RepeatMasker. For chromatin-contact analysis, Hi-C reads were aligned to the final assembly with BWA-MEM2, filtered with Pairtools, and converted into contact matrices using Cooler.

**Fig. S2: Assembly and validation of the Ras3 *D. melanogaster* genome. a,** Chromosome-scale assembly lengths of the Ras3 genome compared with the *D. melanogaster* reference genome dm6.56. **b,** BUSCO assessment of genome completeness for Ras3 and dm6.56, showing the proportions of complete single-copy, complete duplicated, fragmented, and missing BUSCO genes. **c,** Circos representation of the Ras3 genome showing GC content, gene density, and repeat density across chromosomes, calculated in 100-kb windows. **d,** Hi-C contact map of the Ras3 assembly after Knight–Ruiz (KR) normalization, shown at 128-kb resolution, supporting chromosome-scale scaffolding.

**Fig. S3: Chromosome-specific distributions of genomic read counts across parental and hybrid cell lines. a,** Distribution of genomic read counts for each chromosome in the parental Ras3 and ML82 cell lines and the four independently derived hybrid lines HYC1–HYC4. Columns represent cell lines, and rows represent species-resolved chromosomes. D. melanogaster–derived chromosomes are shown in purple, and D. simulans–derived chromosomes are shown in cyan. For each chromosome, the horizontal axis shows the genomic read count, and the vertical axis shows the total genomic span (Mb) associated with each count value. Differences in the position and shape of the distributions reflect chromosome-and species-specific variation in genomic dosage across the hybrid lines. **b,** Proportions of chromosome sequence with and without detectable genomic coverage across the same parental and hybrid cell lines. For each cell line and species-resolved chromosome, stacked bars show the proportion of the chromosome with genomic read counts equal to zero or greater than zero. Orange indicates regions with no detected reads (counts = 0), whereas blue indicates regions with detectable genomic coverage (counts > 0). Percentages are calculated relative to the total analysed span of each chromosome. Large fractions with zero counts indicate extensive loss of the corresponding species-derived chromosome sequence.

**Fig. S4: Convergent loss of the *D. melanogaster Hmr* allele across independently derived hybrid cell lines.** Presence–absence profiles of selected hybrid incompatibility genes and related factors in the parental Ras3 (*D. melanogaster*) and ML82 (*D. simulans*) cell lines and in the four independently derived hybrid lines HYC1–HYC4. The left and right matrices show the *D. melanogaster* and *D. simulans* orthologues, respectively. Blue indicates that the corresponding gene was detected, and red indicates that it was not detected. The *D. melanogaster Hmr* allele was absent from all four hybrid lines, whereas the *D. simulans Hmr* allele was retained. In contrast, the other examined loci showed either consistent retention or hybrid-line-specific patterns of loss, identifying *Hmr* as the only surveyed locus with convergent species-specific loss across all hybrid lines.

**Fig. S5: An interspecies chromosome 3R fusion in hybrid cell line HYC3. a,** Representative HYC3 long reads spanning the fusion junction between *D. melanogaster* chromosome 3R and *D. simulans* chromosome 3R. Each row represents an individual long read with split alignments to the two parental genomes. Purple and cyan segments indicate alignments to *D. melanogaster* and *D. simulans*, respectively, and genomic coordinates are shown above and below the aligned segments. Multiple independent reads spanning the same junction support physical linkage between the two species-derived chromosome segments. **b,** Schematic representation of the inferred chromosome complement of HYC3. Purple and cyan denote *D. melanogaster*– and *D. simulans*–derived chromosomal segments, respectively. The diagram highlights the inferred chimeric chromosome formed by fusion of the *D. melanogaster* 3R segment with the *D. simulans* 3R segment, together with the other retained chromosome arms.

**Fig. S6: Hybrid cell lines retain exclusively *D. melanogaster*–derived mitochondrial DNA.** Species-resolved genomic read coverage across the *D. melanogaster* and *D. simulans* mitochondrial genome contigs. The horizontal axis indicates mitochondrial genome position, and the vertical axis shows log2-transformed genomic read counts plus 1. Orange, red, and purple denote Ras3, ML82, and hybrid cell lines, respectively. Ras3 reads map predominantly to the *D. melanogaster* mitochondrial genome, whereas ML82 reads map predominantly to the *D. simulans* mitochondrial genome. All four hybrid cell lines show coverage only across the *D. melanogaster* mitochondrial genome and no detectable coverage across the *D. simulans* mitochondrial genome, indicating exclusive retention of *D. melanogaster*–derived mtDNA.

**Fig. S7: Allele-specific DNA dosage, RNA abundance, and DNA-normalized transcriptional output across chromosomalorthogroups.** Orthogroups containing corresponding *D. melanogaster* and *D. simulans* genes were identified from protein sequences predicted from the species-resolved long-read transcriptomes. Analyses are shown separately for chromosome arms 2L, 2R, 3L, and 3R and for chromosome X, 4. In the scatterplots, each point represents one orthogroup, with the horizontal and vertical axes showing the *D. melanogaster* and *D. simulans* values, respectively. Point color indicates local point density. **a,** Species-resolved DNA abundance across chromosomal orthogroups in the parental and hybrid cell lines. Gene-level DNA counts were normalized to the total number of sequenced DNA bases in the corresponding sample and rescaled to the mean total number of sequenced DNA bases across samples. **b,** Proportions of orthogroups classified as *D. melanogaster*–biased, unbiased, or *D. simulans*–biased on the basis of normalized DNA abundance. **c,** Species-resolved RNA abundance across the same chromosomal orthogroups. Gene-level RNA counts were normalized to the total RNA count of the corresponding library and rescaled to the mean total RNA count across samples. **d,** Proportions of orthogroups classified as *D. melanogaster*–biased, unbiased, or *D. simulans*–biased on the basis of normalized RNA abundance. **e,** DNA-normalized transcriptional output, calculated separately for each parental allele by dividing normalized RNA abundance by the corresponding normalized DNA abundance. **f,** Proportions of orthogroups classified as *D. melanogaster*–biased, unbiased, or *D. simulans*–biased on the basis of DNA-normalized transcriptional output. For visualization, normalized values were transformed as log2(x + 1), with a pseudocount of 1 added to accommodate zero values. Allelic bias was calculated from the untransformed normalized values as fmel = xmel/(xmel + xsim), where x denotes normalized DNA abundance, normalized RNA abundance, or DNA-normalized transcriptional output, depending on the panel. Orthogroups with fmel > 0.7 were classified as *D. melanogaster–*biased, those with fmel < 0.3 as *D. simulans*–biased, and those with 0.3 ≤ fmel ≤ 0.7 as unbiased. Gray, blue, and orange indicate unbiased, *D. melanogaster*–biased, and *D. simulans*–biased orthogroups, respectively. The reduction in allelic bias after normalization of RNA abundance to DNA dosage indicates that chromosomal transcriptional differences largely reflect underlying allele-specific copy-number differences.

**Fig. S8: Gene Ontology enrichment of *D. melanogaster*-biased transcriptional output in HYC1–HYC4.** Gene Ontology enrichment analysis was performed for *D. melanogaster*-specific orthogroups in HYC1 (**a**), HYC2 (**b**), HYC3 (**c**), and HYC4 (**d**). For each orthogroup, the DNA-normalized RNA abundance of the *D. melanogaster* member was compared with that of the corresponding *D. simulans* member. Orthogroups with log_2_FC > 1 and adjusted *P* < 0.05 were classified as *D. melanogaster*-specific and used as input for enrichment analysis. Dot size indicates the number of input orthogroups assigned to each term, the horizontal axis shows the proportion of input orthogroups associated with the term, and colour indicates the Benjamini–Hochberg-adjusted *P* value.

**Fig. S9: Species-discriminating motifs used for species-resolved quantification of rDNA and X-linked BUSCO genes.** Alignment of the motif sequences used to distinguish *D. melanogaster*-and *D. simulans*-derived reads in motif-scan analyses. Motifs are shown for 18S rDNA, 28S rDNA, and the X-linked BUSCO genes used as normalization references for species-resolved estimation of rDNA copy number. Blue shading marks nucleotide positions that differ between the *D. melanogaster* and *D. simulans* motif sequences and therefore provide species-discriminating information. Gene identifiers are shown on the left, and the corresponding orthogroup identifiers are shown on the right.

**Fig. S10: Relationship between transcript and protein abundance across parental and hybrid cell lines. a,** Relationship between orthogroup-level RNA and protein abundance in Ras3, ML82 and HYC1–HYC4. Each point represents one orthogroup. RNA and protein abundances were log2-transformed before analysis. Solid lines indicate least-squares linear fits. The coefficient of determination (R^2^) is shown for each cell line and indicates the proportion of variation in protein abundance associated with transcript abundance. Point colour indicates local point density. **b,** Pairwise comparison of orthogroup-level protein-to-RNA values across Ras3, ML82 and HYC1– HYC4. Each point represents one orthogroup quantified in both cell lines being compared. The lower triangle shows pairwise scatter plots, with point colour indicating local point density, whereas the upper triangle shows the corresponding coefficients of determination (R^2^). Higher R^2^ values indicate greater conservation of orthogroup-specific protein-to-RNA relationships between cell lines. **c,** Orthogroup-level protein-to-RNA divergence between Ras3 and ML82. Orthogroups are ranked along by log2(*K_p_mel_/K_p_sim_*), *K_p_* difference at each end of the ranked distribution and selected orthogroups are highlighted.

## Supplementary Tables

Table S1. Summary of sequencing datasets generated and analyzed in this study.

Table S2. Transcription quantification generated with IsoQuant from species-resolved long-read transcriptomes.

Table S3. Species-resolved gene-level RNA/DNA ratios.

Table S4. Orthogroups identified with OrthoFinder and used for species-resolved comparative analyses.

## Supplementary Data Files

Data S1. Chromosome-scale genome assembly of the Ras3 *D. melanogaster* cell line.

Data S2. Combined *D. melanogaster–D. simulans* reference genome used for species-resolved analyses of hybrid cell lines.

Data S3. Gene annotation of the combined *D. melanogaster–D. simulans* reference genome.

Data S4. Coding sequences extracted from IsoQuant-reconstructed transcripts across all transcriptome datasets.

Data S5. Predicted protein sequences from the combined *D. melanogaster–D. simulans* reference genome.

