## Supplementary figures and images for "Hybrid *Drosophila* cell lines show selective genome instabilities"

### Supplemental Figure S1

Supplementary Fig. 1

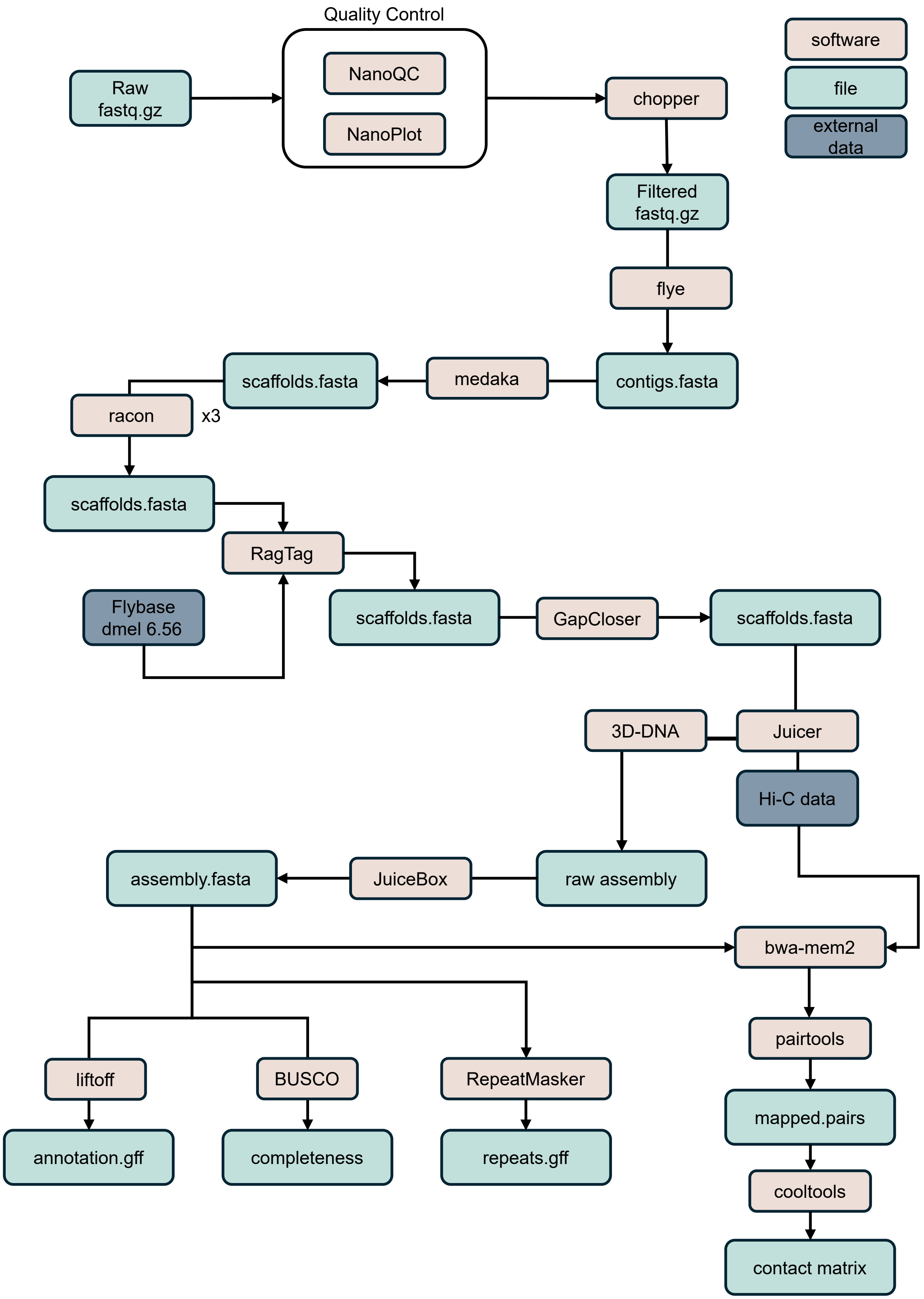

### Supplemental Figure S2

Supplementary Fig. 2

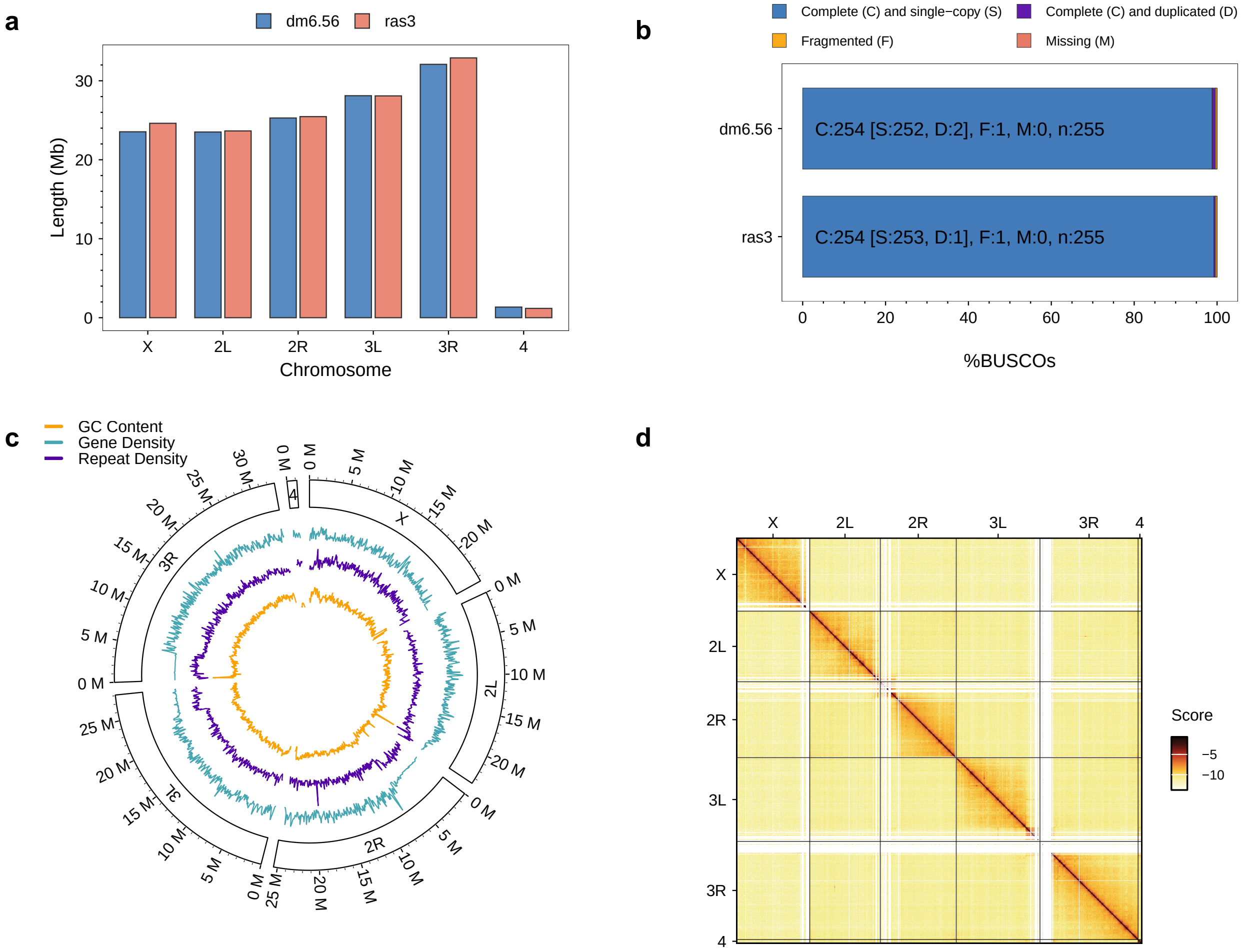

### Supplemental Figure S3

Supplementary Fig. 3

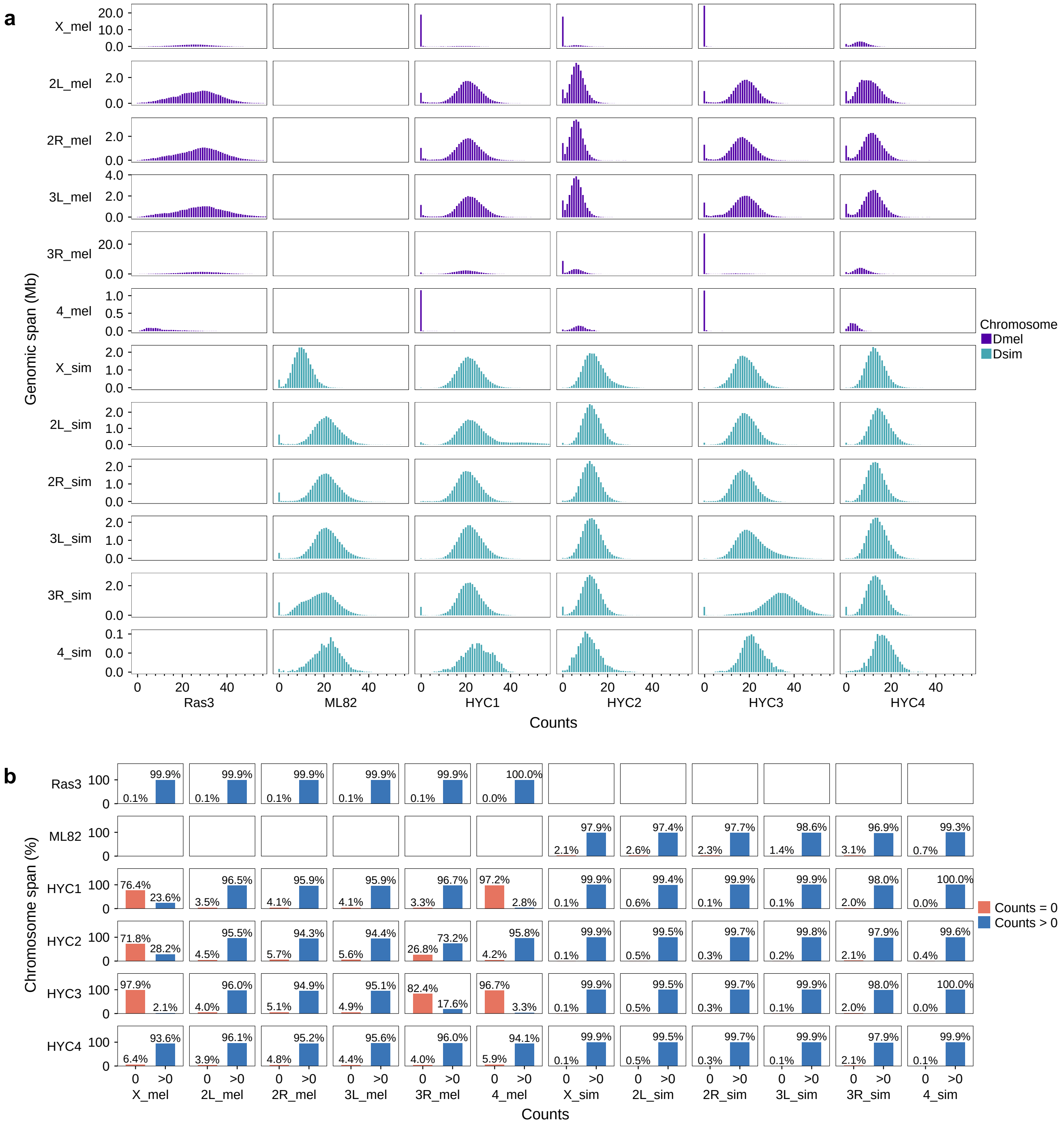

### Supplemental Figure S4

Supplementary Fig. 4

**a**

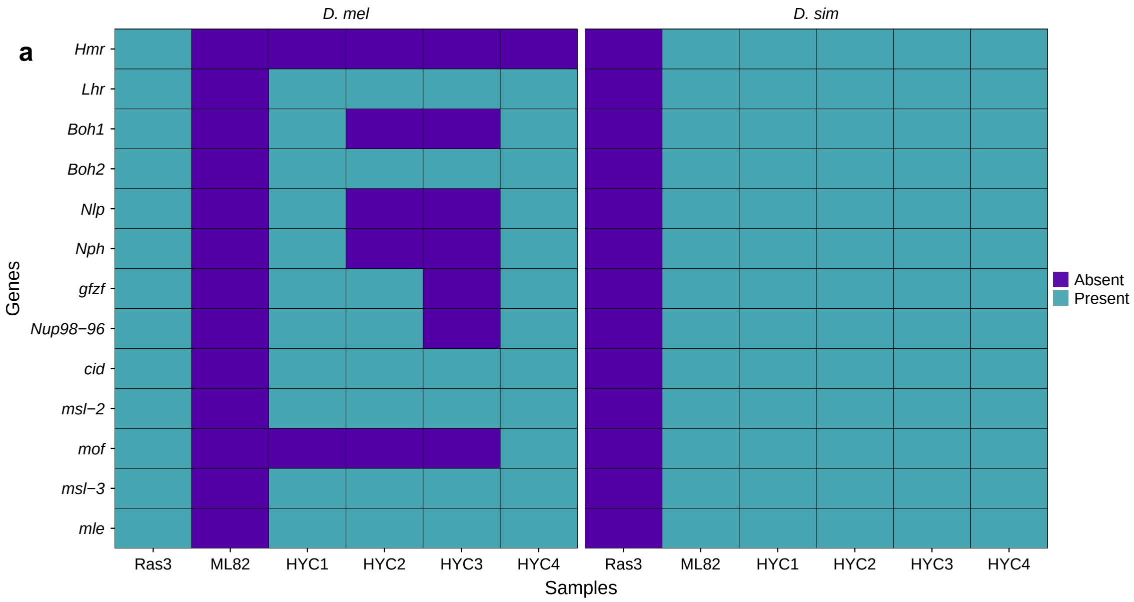

### Supplemental Figure S5

Supplementary Fig. 5

**a**

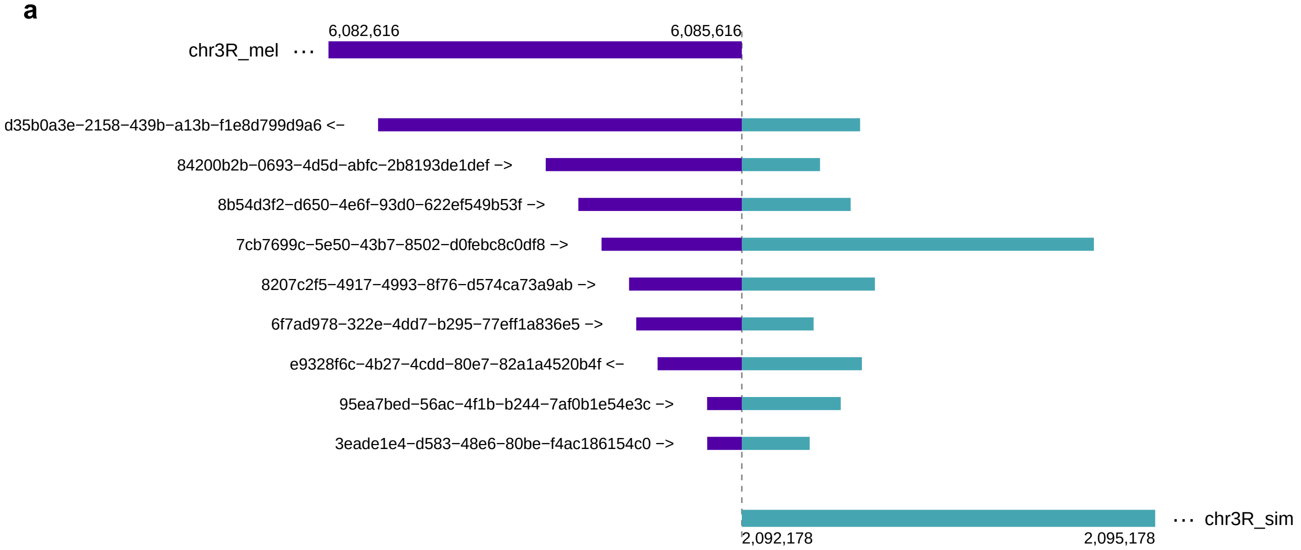


**b**

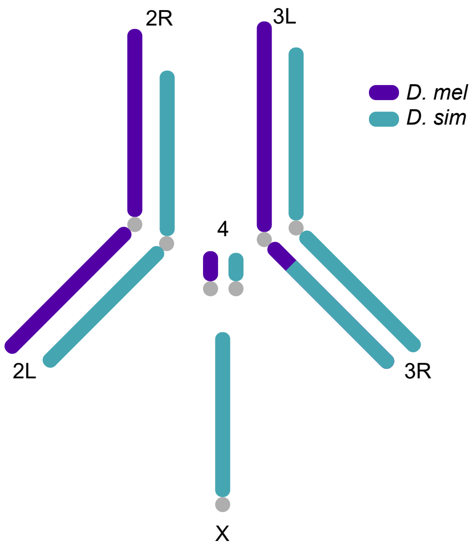

### Supplemental Figure S6

Supplementary Fig. 6

**a**

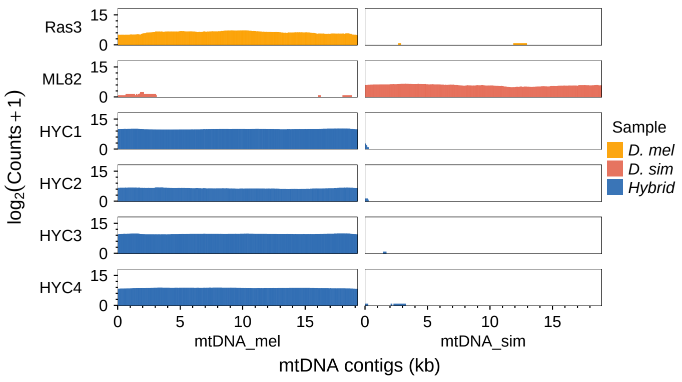

### Supplemental Figure S7

Supplementary Fig. 7

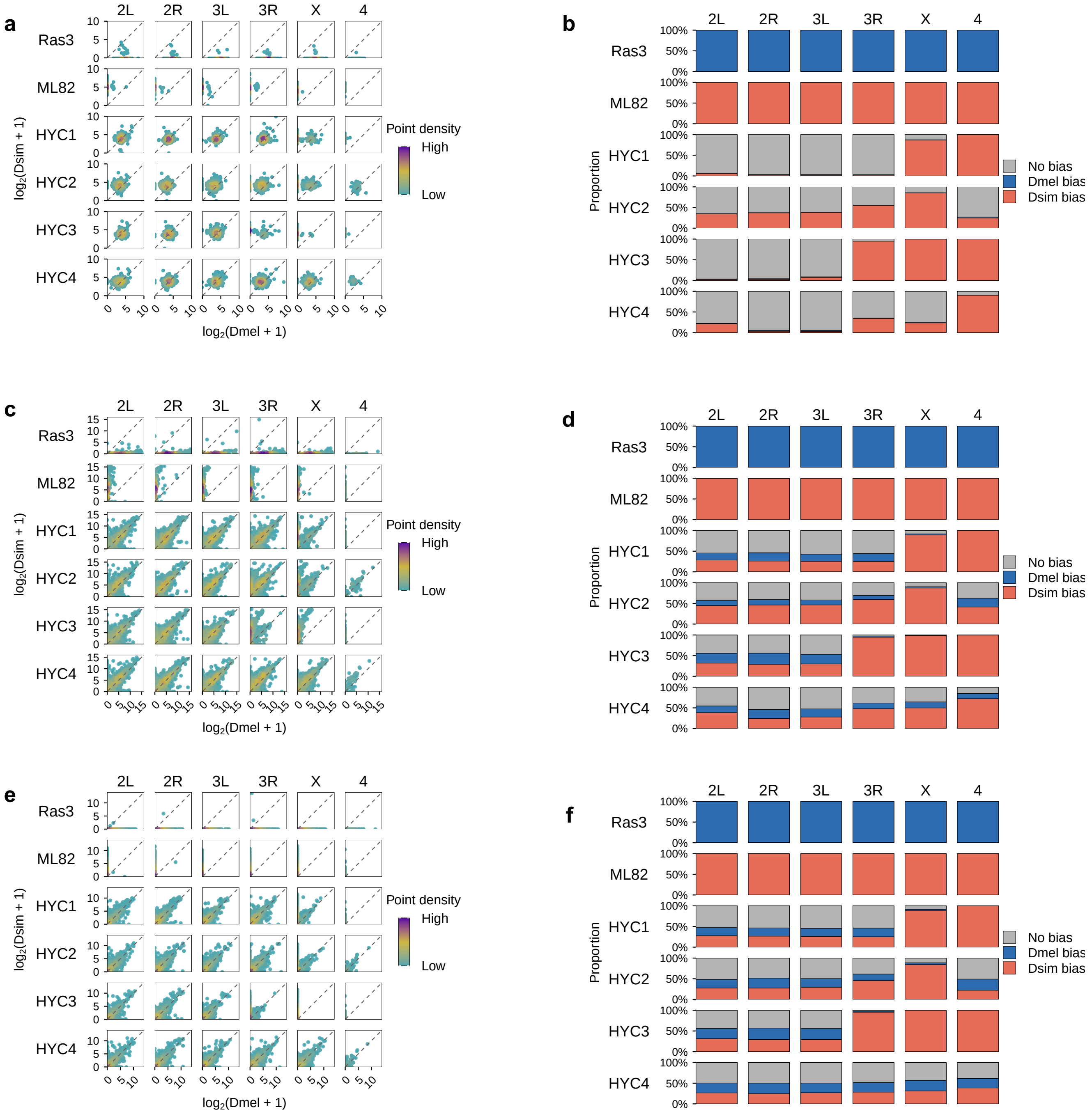

### Supplemental Figure S8

Supplementary Fig. 8

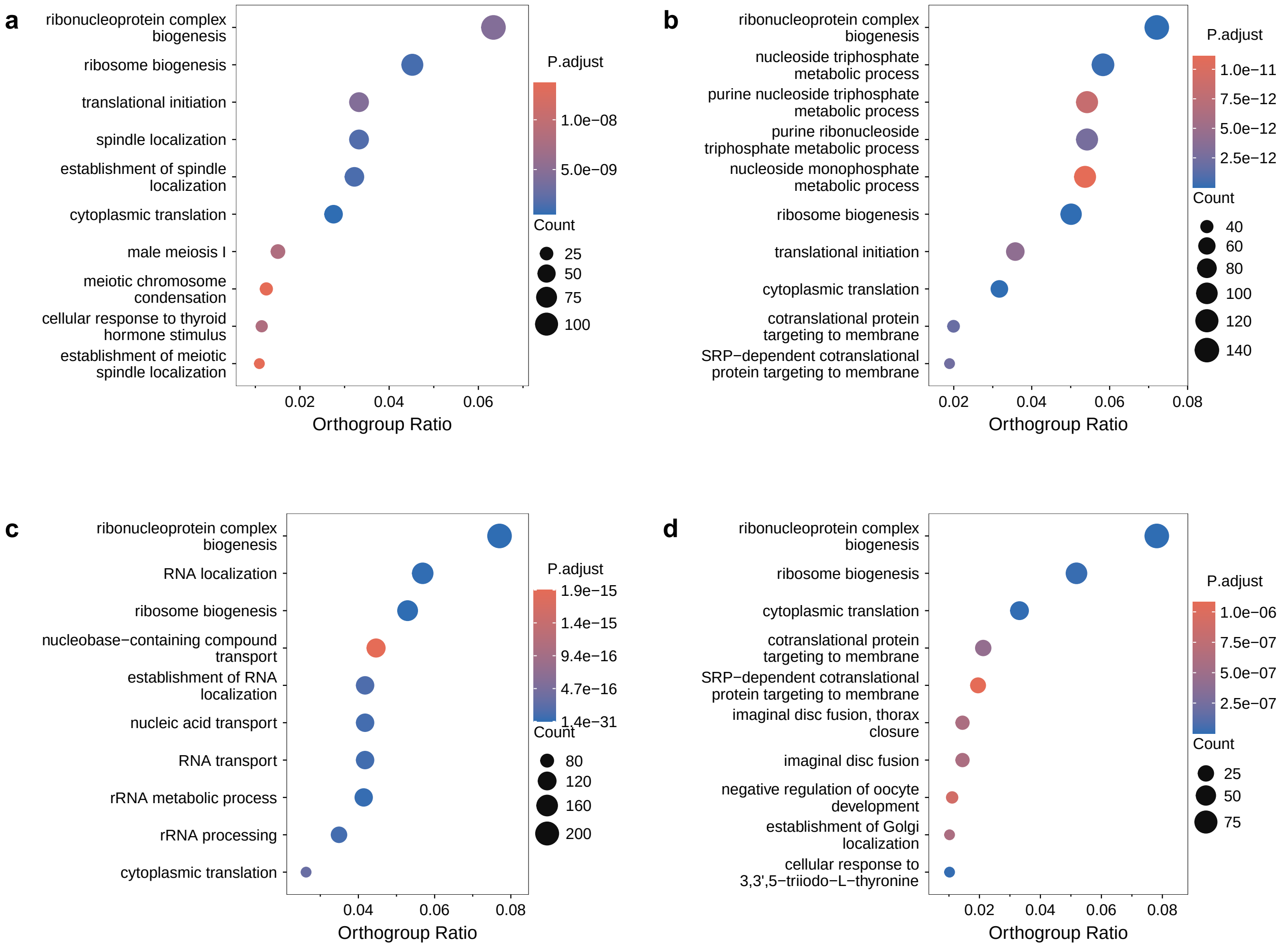

### Supplemental Figure S9

Supplementary Fig. 9

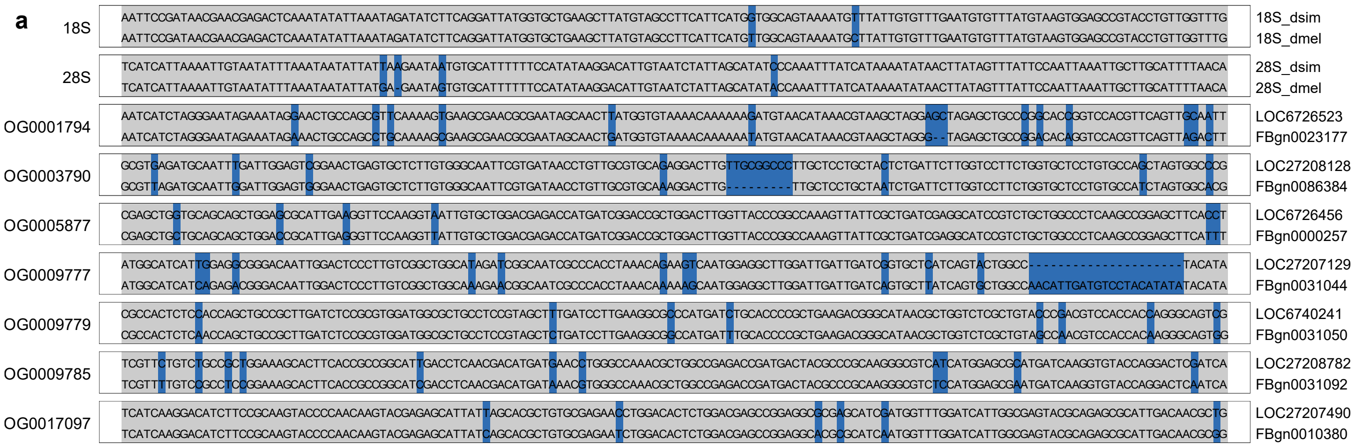

### Supplemental Figure S10

#### Supplementary Fig. 10

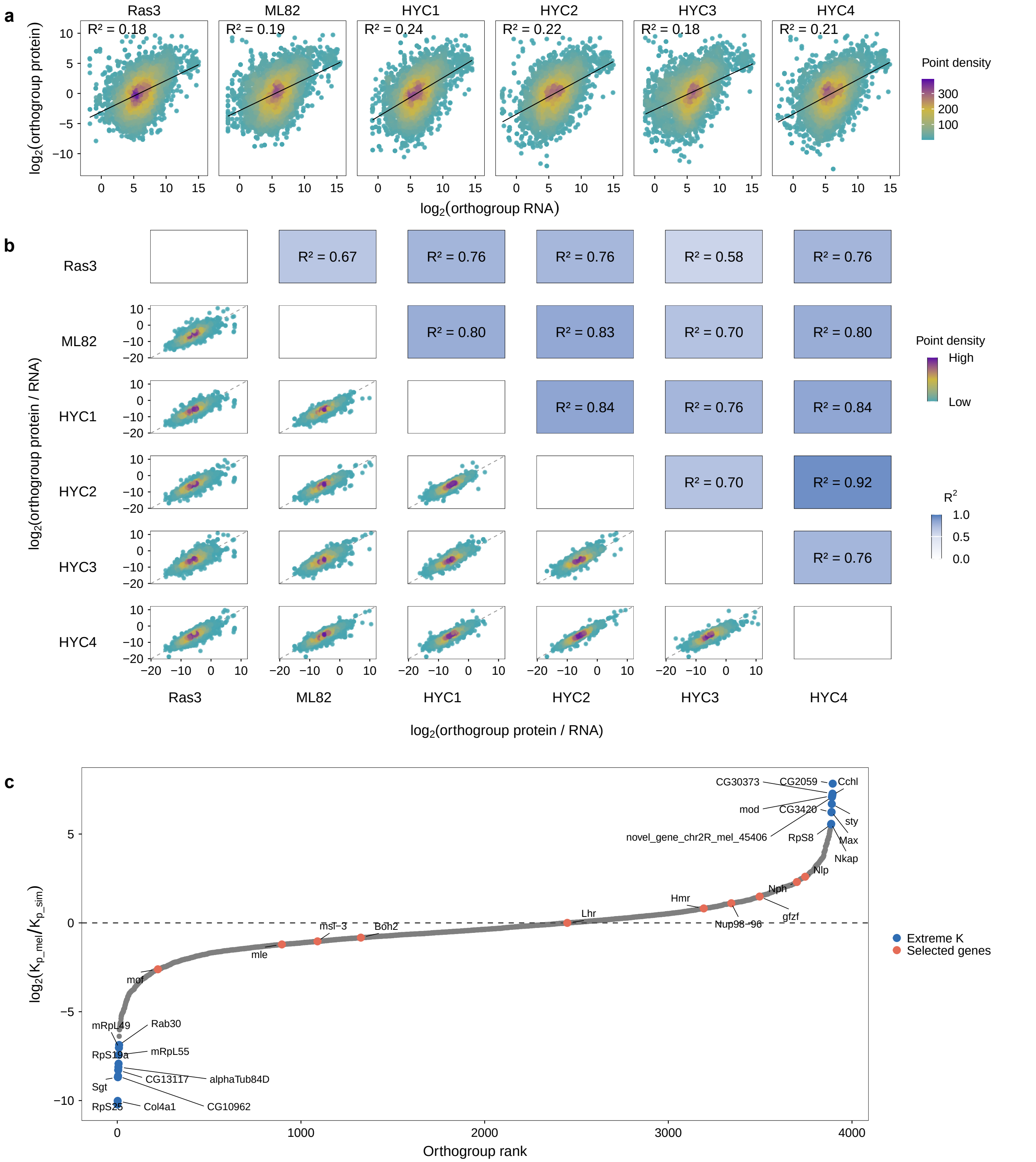
